# HLA-E-VL9 antibodies enhance NK cell and CD8^+^ T cell cytotoxicity against HIV-infected CD4^+^ T cells

**DOI:** 10.1101/2024.07.08.602401

**Authors:** Joyce K. Hwang, Daniel J. Marston, Marina Tuyishime, Hongbing Yang, Daniel Wrapp, Dapeng Li, Simon Brackenridge, McKenzie Frazier, Brianna Rhodes, Caitlin Harris, Max Quastel, A. Brenda Kapingidza, Anna E. Kliszczak, Jacob Gater, Persephone Borrow, Guido Ferrari, Geraldine M. Gillespie, Andrew J. McMichael, Barton F. Haynes, Mihai L. Azoitei

**Affiliations:** Duke Human Vaccine Institute, Duke University School of Medicine, Durham, NC, USA; Department of Medicine, Duke University School of Medicine, Durham, NC, USA; Department of Cell Biology, Duke University School of Medicine, Durham, NC, USA; Department of Surgery, Duke University School of Medicine, Durham, NC, USA; Nuffield Department of Clinical Medicine, University of Oxford, Oxford, UK; Department of Integrative Immunobiology, Duke University School of Medicine, Durham, NC, USA

## Abstract

A major natural killer (NK) cell and CD8^+^ T cell checkpoint is mediated by the inhibitory receptor NKG2A/CD94 and its ligand, HLA-E complexed with 9 amino acid HLA-Ia leader sequence-derived peptides termed VL9 (HLA-E-VL9). Here, we used structure-based design and high throughput library screening to generate antibodies that block NKG2A/CD94 interactions, resulting in direct NK and CD8^+^ T cell cytotoxicity, and trigger NK cell antibody-dependent cellular cytotoxicity (ADCC). Anti-HLA-E-VL9 antibodies limited HLA-E-VL9+ tumor growth in mice, demonstrating checkpoint inhibition activity *in vivo*. HLA-E-VL9 was found to be expressed on HIV-infected cells, and its engagement by HLA-E-VL9 antibodies eliminated infected cells by NK-mediated ADCC. HLA-E-VL9 antibodies also enhanced the killing of HIV-infected cells by NKG2A/CD94^+^ CD8^+^ T cells targeting a novel HLA-E binding HIV Rev-derived epitope. Therefore, anti-HLA-E-VL9 antibodies represent a novel approach to eliminate pathogenic target cells by enhancing NK and CD8+ T cell function and promoting ADCC.

## INTRODUCTION

HLA-E is a non-classical HLA class Ib molecule which, unlike classical HLA class Ia, has limited polymorphism and predominantly presents a 9 amino acid conserved peptide derived from the leader sequence of classical HLA-I and non-classical HLA-G. Endogenous HLA-E-displayed sequence variants center on VM[A/P]PRT[V/I][L/V/I//F]L and are referred to generically as ‘VL9’ [1–4]. HLA-E in complex with VL9 (HLA-E-VL9) binds an inhibitory receptor NKG2A/CD94 that is present on approximately 40% of human natural killer (NK) cells and 5% of circulating CD8^+^ T cells [5] that downmodulates their cytotoxicity [2, 4]. HLA-E also interacts with an activating receptor NKG2C/CD94 [1, 2], but the binding affinity of HLA-E-VL9 peptide complexes for NKG2A/CD94 is greater than that for NKG2C/CD94, such that the inhibitory signal dominates [2, 6, 7]. HLA-E is constitutively expressed at low levels in most normal tissues and at higher levels on leukocytes [8] and in specific tissues such as the placenta, where HLA-E-VL9 interactions with NK NKG2A/CD94 are hypothesized to facilitate tolerance to the fetus [9].

Tumors and viruses exploit the inhibitory function of HLA-E to suppress NKG2A^+^ NK and CD8^+^ T cell killing, making the HLA-E/NKG2A an immune checkpoint of broad relevance. HLA-E is upregulated on diverse tumor types, including cervical and ovarian cancers [10], colorectal cancers [11–13], lung cancers [14], chronic lymphocytic leukemia [15], and circulating tumor cells from patients with pancreatic adenocarcinoma [16]. High levels of HLA-E and/or NKG2A expression within tumors have been correlated with poor disease outcomes [14, 17–20]. Knockdown of HLA-E expression has prevented tumor metastasis in a mouse model of pancreatic adenocarcinoma *in vivo* [16], underscoring the role of HLA-E in immune evasion. A monoclonal antibody against NKG2A, monalizumab, has been developed to enhance NK cell killing of tumor cells [21]. However, when used clinically alone or in combination with other checkpoint inhibitors, efficacy results of NKG2A blockade have been modest for tumor cell control or elimination in humans [22], suggesting that direct blockade of NKG2A alone is insufficient.

Upregulation of HLA-E has also been reported in HIV-1 [23] and is well established in the context of human cytomegalovirus (HCMV) infections [24, 25]. HCMV downregulates classical MHC class Ia to escape T cell detection and encodes its own VL9 peptide within the HCMV-encoded UL40 protein which maintains HLA-E surface expression to evade NK cell attack [24, 25]. Notably, virus-derived peptides bound to HLA-E can also act as important targets for CD8+ T cells. Certain HCMV-encoded VL9 peptide-HLA-E complexes not only bind to NKG2A/CD94, but can also trigger CD8^+^ T cell recognition and drive expansion of antigen-specific T cell responses[24, 26–29]. Other pathogen-derived peptides can also be presented with HLA-E. Hansen et al. demonstrated in a simian immunodeficiency virus (SIV) model that Mamu-E (HLA-E homolog) restricted CD8^+^ T cells elicited by a rhesus CMV vector vaccination mediated SIV control, including viral replication arrest and early clearance of SIV-infected CD4^+^ T cells in >50% of animals [30–33]. Yang et al identified a HLA-E binding HIV-1 Gag peptide and demonstrated priming of HLA-E restricted, HIV-1 specific CD8^+^ T cells *in vitro* [34]. These Gag-specific T cell clones and allogeneic CD8^+^ T cells transduced with these Gag-specific T cell receptors suppressed HIV-1 replication in CD4^+^ T cells *in vitro*. Thus, HLA-E-peptides can function as an immune checkpoint and present endogenous or pathogen peptides to CD8^+^ T cells.

Targeting of upregulated HLA-E-VL9 complexes on target cells by antibodies is an attractive therapeutic strategy because it may simultaneously activate three cytotoxicity mechanisms. NK cytotoxicity may proceed solely through direct interaction of NK cell germline-encoded receptors with ligands for activating NK receptors that have been upregulated and/or ligands for inhibitory NK receptors that have been downregulated on the targets (direct killing) [35]. Thus, first, an antibody binding to HLA-E-VL9 peptide complexes may disrupt the HLA-E/NKG2A checkpoint present on a subset of NKG2A^+^ NK cells, leading to enhanced NK direct killing as has been observed with the anti-NKG2A monoclonal antibody monalizumab [21]. Second, HLA-E-VL9 antibodies may sensitize pathogenic targets by antibody Fab binding and trigger NK mediated target cell killing through Fcγ receptor (FcγR)-engagement via antibody dependent cellular cytotoxicity (ADCC) [36]. Third, an anti-HLA-E-VL9 antibody may bind to NKG2A on a subset of CD8^+^ T cells and enhance TCR-directed CD8^+^ T cell target cell killing.

We previously reported the presence of HLA-E-VL9 antibodies within the natural antibody repertoire of both mice and humans that are predominantly IgM, minimally mutated, and have low affinity for HLA-E-VL9 [37]. Here, we employed structure-based design and high throughput library screening to affinity mature three different anti-HLA-E-VL9 antibodies. Structural analysis revealed the details of their molecular interactions essential for high affinity and specificity. Affinity-matured HLA-E-VL9 antibodies enhanced NK and CD8+ T cell target cell cytotoxicity *in vitro* and in a tumor model *in vivo*. Anti-HLA-E-VL9 antibodies revealed changes in HLA-E-VL9 surface expression upon HIV-infection and eliminated HIV+ CD4+ T cells *in vitro* by mediating NK ADCC and enhancing CD8^+^ T cell activity. Thus, HLA-E-VL9 antibodies represent a promising strategy for eradicating HIV-infected cells and enhancing NK and T cell cytotoxicity for other pathologic cell types, such as tumors with upregulated HLA-E.

## RESULTS

### Engineered anti-HLA-E-VL9 antibodies have high affinity and specificity

Through *in vitro* affinity maturation by yeast surface display, one murine antibody, 3H4v3, achieved high 1:1 binding affinity for HLA-E-VL9 (*K*_D_ = 220 nM) and enhanced direct NK killing of HLA-E-VL9 expressing 293T cells *in vitro* [37]. Here, we further enhanced the affinity of 3H4v3 and of two human IgM anti-HLA-E-VL9 antibodies, CA147 [37] and CA117. Our goals were first to determine if antibody modification could yield monoclonal antibodies (mAbs) with improved affinity or specificity, and second, to investigate whether higher HLA-E-VL9 Ab affinity led to increased enhancement of NKG2A^+^ effector cell cytotoxicity.

The affinities of CA147 and CA117 mAbs were optimized separately by screening scFv libraries that diversified the sequence of the CDR loops [37] (**Figure 1A; Figure S1A, B**). This resulted in the identification of antibodies CA147v24 and CA117v2v8, that bound to HLA-E-VL9 with *K*_D_ values of 2.6 μM and 2.4 μM, respectively (**Figure 1B; Figure S1C**). Compared to the parent antibodies, CA147v24 contained three mutations in the CDRH2 loop (G55R; S56R; T57E), while CA117v2v8 had four mutations in the CDRH1 loop (S32K; S33D; Y34V; Y35M) and three mutations in the CDRL2 loop (N95W; N95aS: S95bK). The increased affinity of CA147v24 and CA117v2v8 enabled us to determine their structure when bound to HLA-E-VL9 **(Table S1)**. Interestingly, 3H4, CA147v24, and CA117v2v8 each interacted differently with HLA-E-VL9 by contacting distinct VL9 amino acid residues (**Figure 1C**). 3H4 primarily interacted with position 1 of VL9, CA147v24 engaged both positions 1 and 5, while CA117v2v8 contacts were centered around position 8 (**Figure 1D**). All antibodies also contacted the HLA-E heavy chain via alpha helices α1 or α2, that form the cleft of the peptide binding pocket, and newly engineered contacts mediated several of these interactions (CA147v24: Arg55, Arg56; CA117v2v8: Lys32; **Figure S1D**). These different binding modes influenced the specificity of the engineered antibodies. From a panel of HLA-E/peptide complexes tested, 3H4 and CA147v24 showed high specificity for HLA-E-VL9 (**Figure 1E**), whereas CA117v2v8 also bound to HLA-E in complex with SARS-CoV-2 and *M. tuberculosis* derived peptides (**Figure S1E**). Sequence analysis of multiple peptides bound by HLA-E revealed that position 8 was mostly conserved to aliphatic residues, which explained the polyreactivity of CA117v2v8 (**Table S2**). Crystallographic analysis of CA117v2v8 bound to HLA-E in complex with *M. tuberculosis* (Mtb) peptide Mtb-44 (RLPAKAPLL) and HIV-1-RL9 (RMYSPTSIL) peptides further supported this analysis, revealing that CA117v2v8 light chain residue W95 closely interacts with the conserved hydrophobic residues at position 8 (**Figure S1F)**. Positions 1 and 5 were more variable, with no peptides containing the same amino acids at both of these two sites, suggesting that CA147v24 may be specific beyond the four peptides tested.

**Figure 1:**
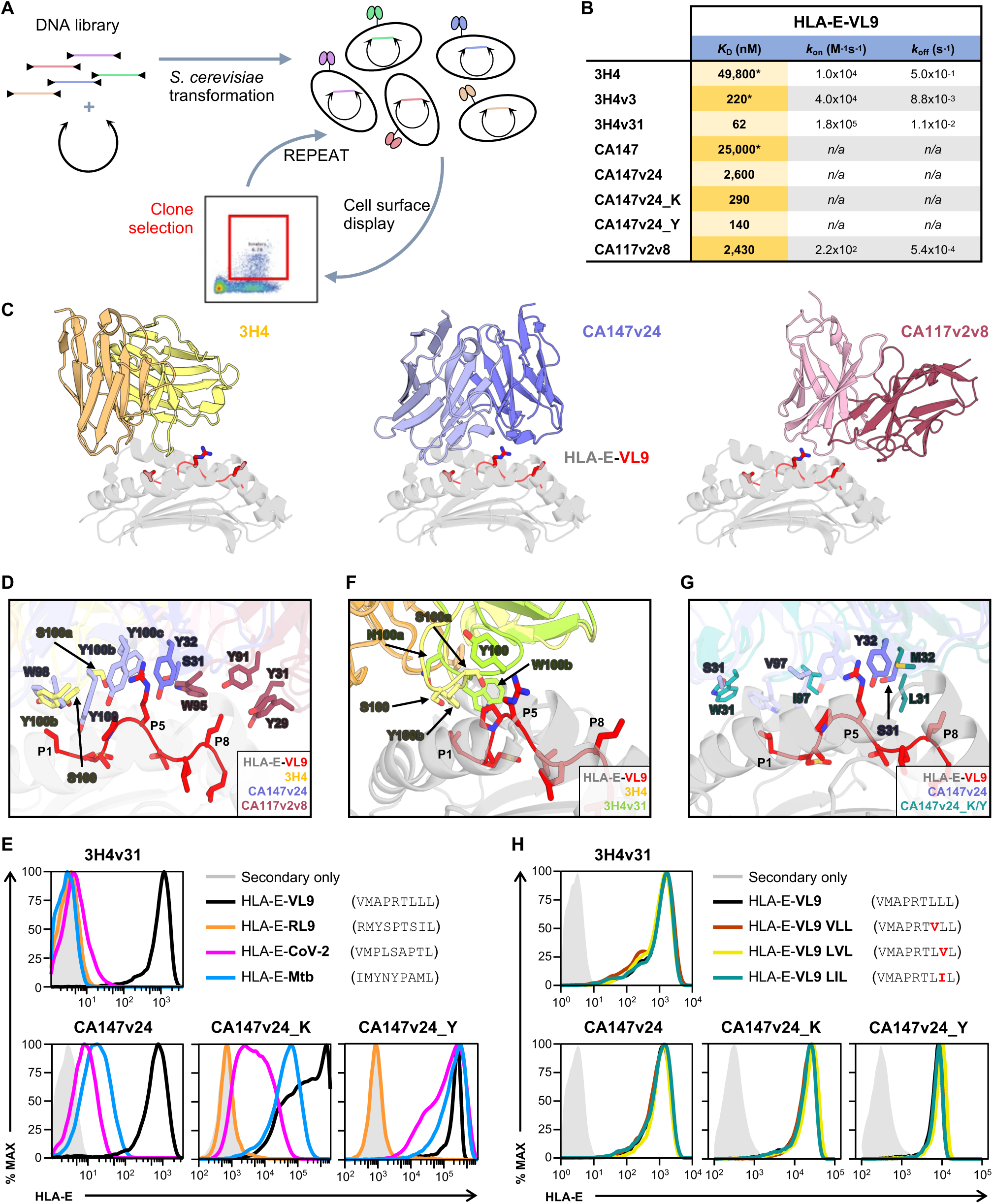
Development and characterization of affinity matured anti-HLA-E-VL9 antibodies. **(A)** Schematic of antibody library screening on the surface of yeast. Mutant scFv plasmid libraries are transfected into yeast and displayed on the cell surface. Clones that enhance HLA-E-VL9 binding are then iteratively enriched through fluorescence activated cell sorting. **(B)** 1:1 binding affinities of anti-HLA-E-VL9 antibodies. * = values from [37] **(C)** Crystal structures of three different antibodies in complex with HLA-E-VL9. HLA-E heavy chain is shown in *gray*, and the VL9 peptide is in *red*. **(D)** Detailed view of the interactions between the antibodies in (C) and the VL9 peptide. Antibodies colored as in (C). Peptide positions of interest are labeled P1, P5 and P8. Interacting antibody residues are labeled and shown in sticks. **(E)** FACS binding of anti-HLA-E-VL9 antibodies to HLA-E in complex with different peptides derived from HIV (RL9), SARS-CoV-2 or *M. tuberculosis* (Mtb). **(F)** Interface view of the crystal structure of 3H4v31 (*green*) aligned on the previously determined complex of 3H4 bound to HLA-E VL9. 3H4v3 and HLA-E-VL9 are colored as in (C). Residues mutated between 3H4v3 and 3H4v31 are numbered and shown in sticks. **(G)** Models of CA147v24_K and Y in complex with HLA-E-VL9 overlayed onto the CA147v24 + HLA-E-VL9 crystal structure colored as in (C). Mutated residues in CA147v24_K and CA147v24_Y are numbered and shown in sticks. **(H)** FACS binding of anti-HLA-E-VL9 antibodies to HLA-E in complex with different VL9 isoforms.

With high-resolution structures now available (**Table S1**), we further improved the affinity of 3H4v3 and CA147v24 by building directed libraries (**Figure S2A-B**). Following library selection, one clone, named 3H4v31, contained three additional mutations relative to the 3H4v3 template (S100aR, Y100bW, and G100cN). The recombinantly expressed 3H4v31 IgG1 antibody bound to HLA-E VL9 with an affinity of 11.5 nM, representing a 19-fold increase over 3H4v3 and a ∼4300-fold increase compared to the originally isolated 3H4 [37] (**Figure 1B; Figure S2C)**. The crystal structure of the unbound 3H4v31 Fab confirmed that the affinity maturation process altered the CDRH3 loop while maintaining the overall structure of the 3H4 Fab (**Figure 1F**). Alignment of 3H4v31 to the previously determined co-crystal structure of wildtype 3H4 (3H4 WT) bound to HLA-E-VL9 (PDB ID: 7BH8) revealed that the affinity-matured CDRH3 loop of 3H4v31 was positioned across the VL9 peptide and contacted Pro4 and Arg5 of the peptide (**Figure 1F**).

Affinity maturation of CA147v24 by yeast display (**Figure S2B**) identified two new versions, CA147v24_K and CA147v24_Y (**Figure S2C, D**) that bound to purified HLA-E-VL9 with a *K*_D_ of 290 nM, representing a 9-fold increase over CA147v24 and a ∼90-fold increase compared to the natural CA147 as IgG [37] **(Figure 1B)**. CA147v24_Y, which had one additional mutation compared to CA147v24_K (CDRH1: S31W), had a twofold higher affinity.

To gain insight into the interaction modes of these antibodies, we generated models of their complex with HLA-E-VL9 using AlphaFold3 (**Figure 1G**). This was deemed suitable, since AlphaFold3 correctly predicted the crystal structure complex of CA147v24 and HLA-E-VL9 (**Figure S2E**). Changes at positions 31 and 32 enabled the new Leu31 residue, present in both CA147v24_K and CA147v24_Y, to interact with position 8 of the peptide. These changes led to a loss of a degree of specificity for CA147v24_K and CA147v24_Y compared to CA147v24, based on measured interactions with HLA-E complexed with *M. tuberculosis* or SARS-CoV-2 derived peptides (**Figure 1E**). Due to natural leader sequence variation, multiple VL9 isoforms can be displayed on HLA-E, therefore we tested antibody binding to complexes containing diverse VL9 peptides (**Figure 1H**). Both 3H4 and CA147 derived antibodies bound tightly to all the complexes tested, revealing their broad recognition of HLA-E-VL9. Thus, our results illustrate the development of multiple antibodies that bind to HLA-E-VL9 with high affinity. Structural and biochemical analyses revealed that these antibodies employ different interaction modes that modulate their specificity.

### Anti-HLA-E mAbs enhance NK mediated natural cytotoxicity and ADCC *in vitro*

We next assayed the ability of anti-HLA-E-VL9 mAbs to enhance direct cytotoxicity by measuring the killing of a K562 human leukemia cell line [38] that had been modified to express HLA-E-VL9 as a single chain trimer of HLA-E, beta-2-microglobulin and VL9 peptide (K562^HLA-E-VL9^) [34]. Effector cells in these assays were the NKG2A+ NK cell line, NK-92, that lacks CD16/FcγRIIIa (ATCC CRL-2407) [39]. NK-92 cytotoxicity, measured by 51-chromium (^51^Cr) release assays [40], was significantly decreased against K562^HLA-E-VL9^ targets compared to K562 cells lacking HLA-E-VL9, demonstrating the suppression of HLA-E-VL9 on killing by NKG2A-expressing NK-92 cells (**Figure 2A**). In this setting, the addition of 3H4v31 IgG1 enhanced NK-92 killing of K562^HLA-E-VL9^ by 1.5 to 2.2-fold, over a range of concentrations between 0.016 μg/mL to 10 μg/mL, with a dose dependent relationship (Kendall Tau correlation = 0.5450, P = 0.0004) (**Figure 2B**). Similar effects were not observed with the low affinity 3H4 IgG1 (**Figure S3A**), demonstrating that enhancement of NK cytotoxicity was modulated by the affinity for HLA-E-VL9. 3H4v31 IgG1 had no intrinsic cytotoxicity, as no killing occurred when this mAb was added to the K562^HLA-E-VL9^ cell culture in the absence of NK-92 NK cells (**Figure S3B**). Both human-derived CA147v24_K and CA147v24_Y affinity matured HLA-E-VL9 mAbs also enhanced direct NK cell killing, by 1.9-fold and 1.8-fold over isotype control, respectively. (**Figure 2C**). Thus, high-affinity anti-HLA-E-VL9 mAbs enhanced NK cell-mediated direct cytotoxicity of K562^HLA-E-VL9^ targets.

**Figure 2:**
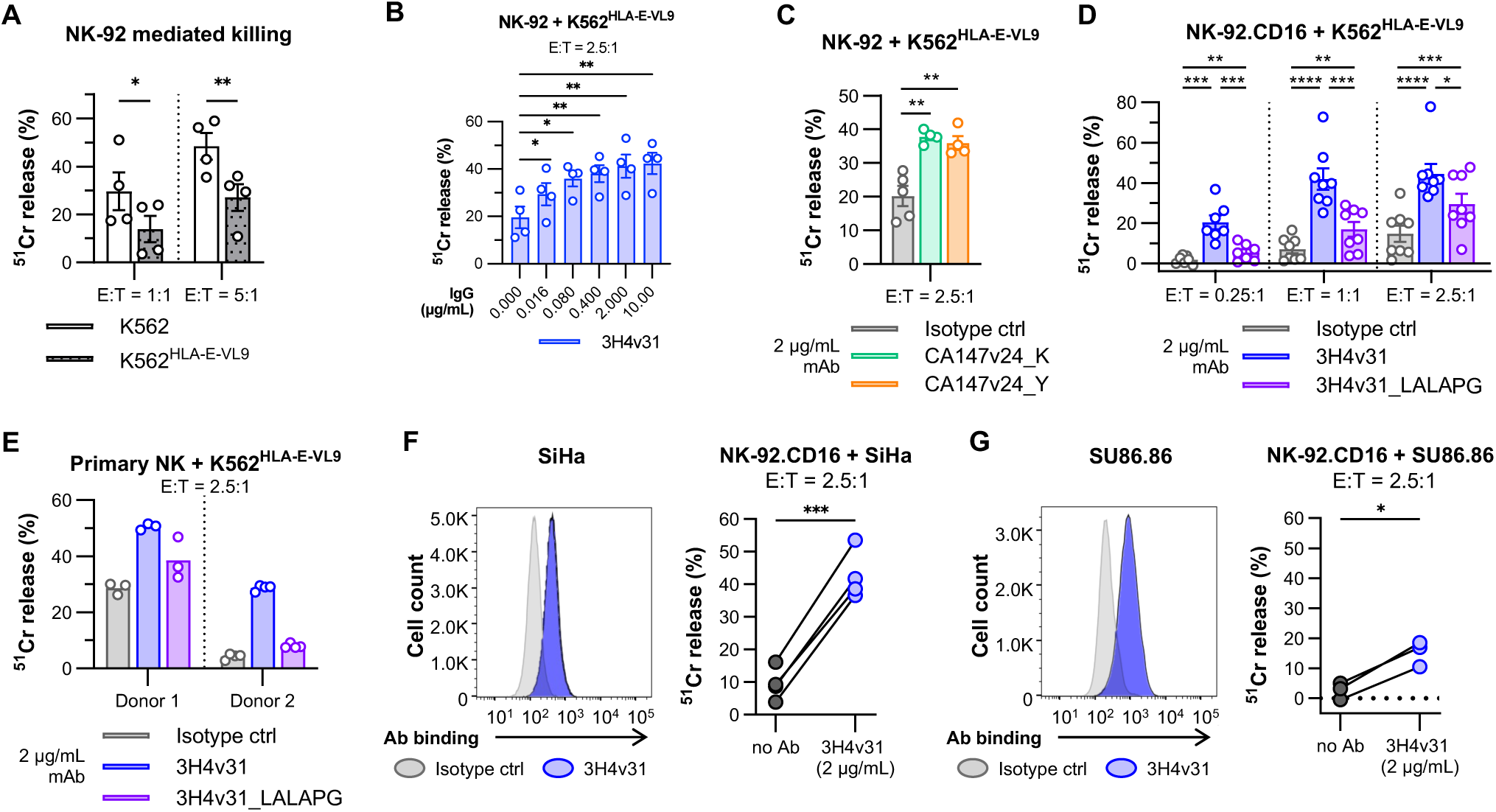
HLA-E-VL9 mAbs enhance target cell killing by NK cells. **(A)** Target cell killing of K562 or K562^HLA-E-VL9^ target cells by NKG2A^+^ NK cell line NK-92 as assessed by ^51^chromium (^51^Cr) release assay. n = 4 independent experiments. **(B)** Target cell killing by NK-92 with increasing concentration of affinity-matured 3H4v31 IgG1 (3H4v31). n = 4 independent experiments. **(C)** Target cell killing by NK-92 in the presence of 2 μg/mL of mAbs CA147v24_K and _Y. n=4-5 independent experiments. **(D)** Target cell killing by NK-92.CD16 NK cells in the presence of 2 μg/mL of 3H4v31, 3H4v31 with LALAPG Fc mutation (3H4v31_LALAPG), and isotype control. n = 7-8 independent experiments. **(E)** Target cell killing by primary NK cells isolated from two human donors in the presence of 2 μg/mL antibody at E:T ratio of 2.5:1. Dots indicate well replicates. **(F-G)** 3H4v31 binding to SiHa, n = 4 independent experiments (F), and SU86.86, n = 3 independent experiments (G). *Left*: FACS binding with 3H4v31 and isotype control. Histograms are representative of at least 2 repeats. *Right*: Target cell killing by NK-92.CD16 cells as measured by ^51^Cr assay. Error bars represent standard error of the mean. Statistical analysis was performed with two-sample t-test (C). *p < 0.05, **p < 0.01, ***p < 0.001, ****p < 0.0001.

We next asked whether 3H4v31 could also mediate NK cell ADCC. Here, we used as effectors NK-92 transduced with CD16 (NK-92.CD16) [41, 42] and primary PBMC NK cells. Additionally, we constructed a “LALAPG” Fc variant of 3H4v31, in which FcγR-dependent function was abolished (3H4v31_LALAPG) [43, 44] (**Figure S3C**), capable of mediating HLA-E-VL9/NKG2A blockade but not ADCC, enabling us to discriminate the relative contributions of ADCC-dependent and ADCC-independent mechanisms of NK enhancement. 3H4v31_LALAPG mediated 20%-50% (depending on the E:T ratio) of the killing enhanced by 3H4v31 of K562^HLA-E-VL9^ cells by NK-92.CD16 **(Figure 2D)**, and 15%-45% (depending on the donor) of the killing enhanced by 3H4v31 of K562^HLA-E-VL9^ cells by primary NK cells **(Figure 2E)**. These results were consistent with anti-HLA-E-VL9 antibodies mediating two distinct NK cytotoxicity mechanisms: enhancing NK direct killing through HLA-E-VL9/NKG2A blockade and killing mediated by CD16 FcR-mediated ADCC.

In addition to directly measuring target cell death via ^51^Cr release, we also assessed the upregulation of lysosome associated membrane protein-1 (CD107a) on NK cells in the presence of HLA-E-VL9 specific mAbs as a marker of NK cell degranulation. NK-92.CD16 cells exhibited 4.4-fold, 5.2-fold, and 4.4-fold greater CD107a+ surface expression upon co-culture with K562^HLA-E-VL9^ in the presence of mAbs 3H4v31, CA147v24_K, or CA147v24_Y, respectively, compared to isotype control (**Figure S3D**). Only a minor increase in CD107a^+^ NK cells compared to isotype control was observed on NK-92.CD16 cells in the ADCC-independent conditions using the 3H4v31_LALAPG antibody (**Figure S3D**) or NK-92 effectors with 3H4v31 antibody (**Figure S3E**), demonstrating that only ADCC-mediated activation produced sufficient effector activation to detect degranulating cells. Primary circulating NK cells isolated from peripheral blood mononuclear cells (PBMC) of healthy donors also demonstrated increased frequency of CD107a positivity upon co-culture with K562^HLA-E-VL9^ in the presence of mAb 3H4v31 (3.1-fold greater than with isotype control mAb) **(Figure S3F).**

We next tested the ability of 3H4v31 to bind and enhance killing of targets expressing endogenously processed peptide that is presented by HLA-E-VL9 on the cell surface. To this end, we used 3H4v31 to identify cell lines expressing HLA-E-VL9, including SiHa (a cervical squamous cell carcinoma cell line) (**Figure 2F**) and SU86.86 (a pancreatic ductal adenocarcinoma cell line) (**Figure 2G**). In ^51^Cr release assays, the killing of these malignant tumor cell line targets by NK-92.CD16 was also enhanced (**Figure 2F, G**).

These results demonstrated that human and mouse anti-HLA-E-VL9 mAbs enhanced both direct killing and ADCC *in vitro* by NKG2A+ NK cell lines and by PBMC NK cells. Importantly, these antibodies recognized both transfected single chain trimer and endogenously expressed forms of HLA-E-VL9 on the surface of HLA-E-VL9^+^ targets.

### Anti-HLA-E-VL9 mAb enhance NK HLA-E-VL9^+^ tumor control *in vivo*

We next asked if the observed anti-HLA-E-VL9 mAb-mediated enhancement of NK killing of K562^HLA-E-VL9^ cells *in vitro* could translate into an anti-tumor effect *in vivo.* We used immunodeficient NOD/SCID/IL2rγ^null^ (NSG) mice that lack endogenous B, T, or NK cells (**Figure 3A**). K562^HLA-E-VL9^ cells were implanted subcutaneously into the right flank of the mice, which grew into palpable tumors 1-2 weeks later. Mice were treated on days 10, 14, 16, and 18 following cell implantation with intra-tumoral injection of NK-92 cells and either 3H4v31 or control mAb (**Figure 3B**). An additional group received control mAb but no NK-92 cells. IL-2 was provided with all NK cell and/or mAb treatments to enhance NK-92 viability. The tumor growth in mice treated with control mAb and NK-92 was similar to that of mice treated with control mAb alone (without NK-92 cells), consistent with suppression of NK cell activity by the K562^HLA-E-VL9^ tumor (**Figure 3B**). Tumor growth was slower in mice receiving NK-92 cells mixed with 3H4v31 mAb (**Figure 3B**). The average tumor growth rate was 96 mm^3^ per day from days 10 to 18 in the 3H4v31 treatment group, compared to 363 mm^3^ per day for the NK-92 cells with control mAb group (mixed effect model, p = 0.00006) and 342 mm^3^ per day for the control mAb without NK cells group (mixed effect model, p = 0.0000008) (**Figure 3B**). Tumor growth was significantly slower in the group treated with 3H4v31 and NK-92 cells compared to control mAb and NK-92 on two additional replicates: 170mm^3^ per day during days of treatment in the 3H4v31 treatment group compared to 372 mm^3^ per day for the control mAb (mixed effect model, p = 0.0212) (**Figure 3C**), and 33 mm^3^ per day during in the 3H4v31 treatment group compared to 140 mm3 per day for the control mAb (mixed effect model, p = 0.0010) (**Figure 3D**).

**Figure 3:**
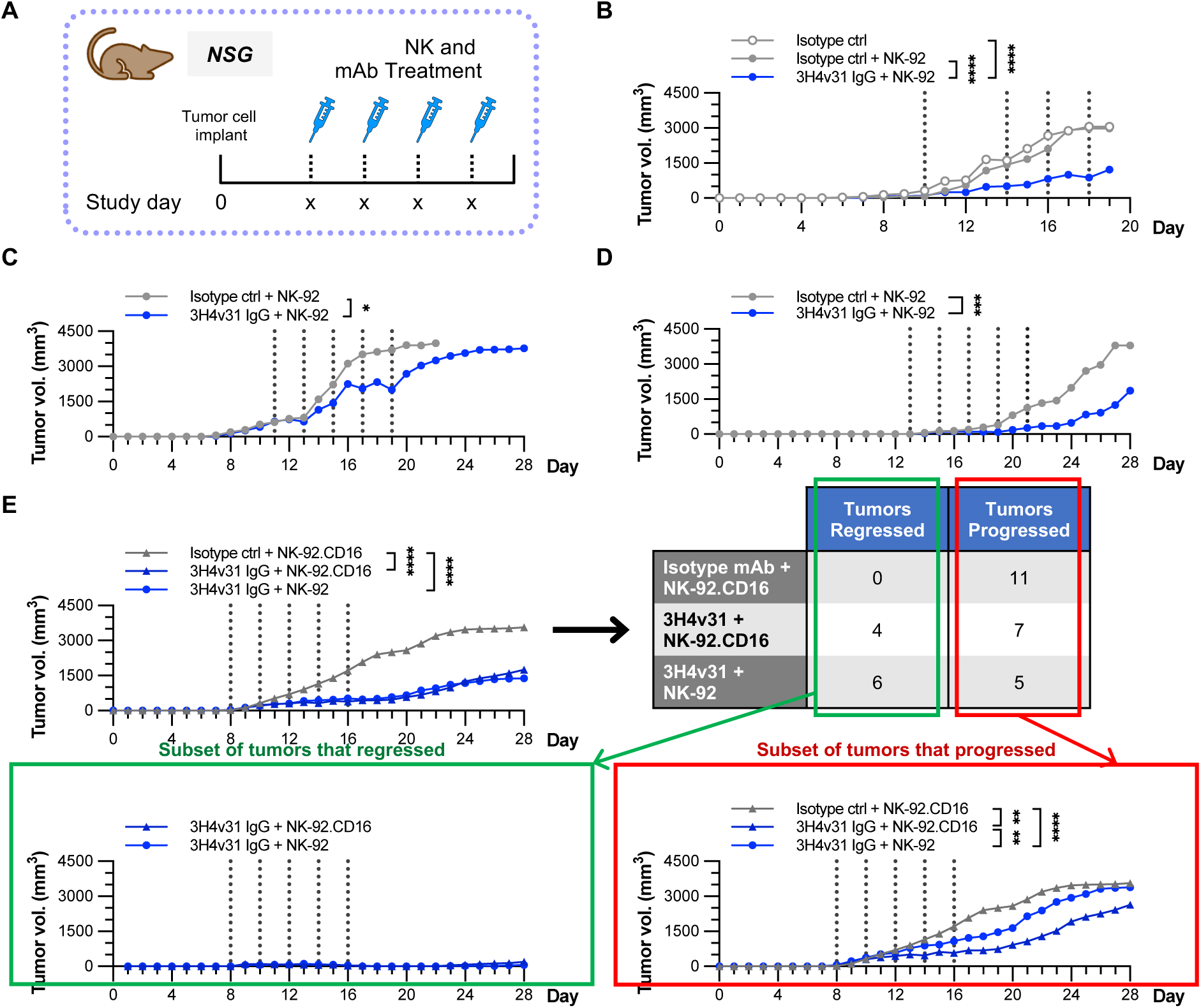
HLA-E-VL9 mAbs retard HLA-E-VL9 expressing tumor growth *in vivo.* **(A)** Nod scid gamma mice were implanted subcutaneously with K562^HLA-E-VL9^ cells. On the days indicated (*vertical dotted lines*), mice received treatments comprised of intratumoral injection of mixtures of 10 million NK effectors and/or 990 μg of mAbs. **(B-E)** Average tumor size per day is plotted for each indicated treatment group**: (B)** n = 5 mice per treatment group. **(C)** n = 10 mice per treatment group. **(D)** n = 11 mice per treatment group. **(E)** n = 11 mice per treatment group. Table lists numbers of mice in each group with tumors that regressed to non-palpable size on treatment or did not grow (“Tumors regressed”) or grew (“Tumors progressed”) throughout treatment. *Green* and *red* boxes show average growth curves of tumors that regressed and tumors that progressed respectively. Statistical analysis was performed using mixed effects models of tumor growth rate. Asterisks indicate statistical significance. *p < 0.05, ***p < 0.001, **** p < 0.0001.

To test if 3H4v31 might mediate greater enhancement of NK tumor killing with the addition of ADCC-competent effectors, we treated NSG mice engrafted with K562^HLA-E-VL9^ tumors with 3H4v31 and either NK-92.CD16 cells or NK-92 cells (**Figure 3E**). An additional group received control mAb and NK92.CD16 cells. In this experiment, tumors were treated earlier post-tumor implantation, on days 8, 10, 12, 14, 16. Several tumors in the two groups treated with 3H4v31 had no measurable growth or regressed on treatment to the point of being non-palpable **(Figure S4A**): 4 of 11 tumors in the NK92.CD16 treated group, and 6 of 11 tumors in the NK-92 treated group regressed (**Figure 3E – *table*).** By comparison, no tumors regressed in the control mAb treated group (**Figure 3E – *table;* Figure S4A**). A difference in tumor growth rate between the groups receiving NK-92.CD16 cells with 3H4v31 versus NK-92 cells with 3H4v31 were masked by the subset of tumors that had regressed, as the average tumor volume was very low in this subset of both groups (**Figure 3E - *green box***). Among the subset of tumors that progressed (**Figure 3E - *red box***) the tumor growth rates were slower in the mice that received NK-92.CD16 with 3H4v31 (rate 68.9 mm^3^/day between days 8-16) compared to the mice that received NK-92 with 3H4v31 (rate 128 mm^3^/day between days 8-16) (mixed effect model, P = 0.0092). Thus, ADCC-incompetent effectors (NK-92) and ADCC-competent effectors (NK-92.CD16) both mediated potent control of K562^HLA-E-VL9^ tumors.

To visualize the cellular interactions underlying the anti-tumor effect of 3H4v31 in the presence of NK cells, tumors from mice treated for 28 hours with one dose of NK-92 cells plus 3H4v31 or NK-92 cells with control antibody were cryopreserved and five-micron sections were formaldehyde-fixed and incubated with anti-CD56 or caspase 3 antibodies to detect NK cell presence and anti-cleaved caspase 3 to detect apoptotic cell death. Tumor sections from both groups of mice demonstrated the presence of NK-92 cells (Figure S4B, C). Apoptosis was detected at higher levels in the 3H4v31-treated tumors compared to the CH65-treated tumors consistent with anti-HLA-E-VL9-specific enhancement of NK mediated cytotoxicity (Figure S4B, C). NK-92 cells were also specifically detected in contact with apoptotic K562^HLA-E-VL9^ cells in the 3H4v31-treated tumors but not in the control mAb-treated tumors (Figure S4B).

### HLA-E-VL9 is expressed on the surface of HIV-infected primary CD4+ T cells

HLA-E is of renewed importance to develop novel strategies to eliminate HIV, as it has been demonstrated that T cells restricted by Mamu-E (the rhesus orthologue of HLA-E) elicited by CMV vector vaccination enabled ∼55% of macaques to clear infection following SIV challenge [33]. Moreover, HLA-E specific T cells have been shown to suppress HIV-1 replication in infected CD4+ T cells *in vitro* [34]. However, until now, high affinity HLA-E-VL9 peptide specific mAbs have not been available to define the expression of HLA-E-VL9 co-expressed on HIV-infected cells, and so the possibility that HLA-E-VL9 co-expression may thwart NKG2A+ killing by NKG2A/CD94-expressing T cells has not been tested. Additionally, while some have reported increased expression of HLA-E on HIV-infected CD4+ T cells [23], others have conversely reported that the Nef proteins of some primary HIV strains decrease HLA-E surface expression [45]. Still others have proposed that HIV Nef affects HLA-A and -B but does not affect HLA-E surface expression [46]. While this prior work raises the possibility that HLA-E-peptide complexes may represent a therapeutic target on HIV-infected cells, these studies relied on the use of antibodies that detected both peptide-bound, as well as peptide-free forms of HLA-E [47]. Thus, the surface expression of HLA-E-VL9 on HIV-infected cells has not been directly measured.

To assay HLA-E-VL9 expression on HIV-infected cells, we used 3H4v31 to conduct a time-course experiment assaying HLA-E-VL9 surface expression on primary CD4+ T cells isolated from two donors. Binding was assayed on CD4 cells either before or following activation for 72 hours (to permit infection with the HIV infectious molecular clone, WITO) then at various timepoints over a further 72 hour period after infection [48]. We developed a 3H4v31 mAb with a murine IgG kappa Fc matching the commercially available anti-HLA-A2 clone BB7.2, enabling concurrent analysis of HLA-A2 levels on HLA-A2 donor CD4+ T cells. To control for any virus-specific effects on HLA-E-VL9 expression, we also examined CD4+ T cells that were also activated and cultured for a further 72 hours in parallel with the HIV-infected cells, but not HIV-infected (“mock-infected”). We separately assayed the p24+ CD4+ subset of infected cells, and the p24+ CD4- subset of infected cells representing infected cells at later stages of the viral lifecycle where Nef-mediated CD4 downregulation had occurred **(Figure S5A)**.

We observed that HLA-E-VL9 expression was increased on activated CD4^+^ T cells by 2.7-fold at 48 hours of mock-infection on donors 1 and 2, and by 3.2-fold on donor 1 and 2.4-fold on donor 2 at 72 hours of mock-infection compared to on resting cells (**Figure 4A**). The p24^+^ CD4^+^ subset of WITO HIV-1 infected cells also expressed elevated levels of HLA-E-VL9, 2.5-fold on donor 1 and 2.6-fold on donor 2 at 48 hours post-infection, and 3.2-fold on donor 1 and 2.4-fold on donor 2 at 72 hours post-infection, similar to mock-infected cells (**Figure 4A**). These data suggest that HLA-E-VL9 upregulation is a function of CD4^+^ T cell activation rather than a virus-specific effect. However, the p24^+^ CD4^-^ subset of infected cells expressed lower surface levels of HLA-E-VL9 than the activated uninfected or p24^+^ CD4^+^ subset of infected cells (**Figure 4A**), consistent with an effect concurrent with the downregulation of CD4 and HLA-A2 by HIV-1 Nef **(Figure 4B, Figure S5B)** [49–51].

**Figure 4:**
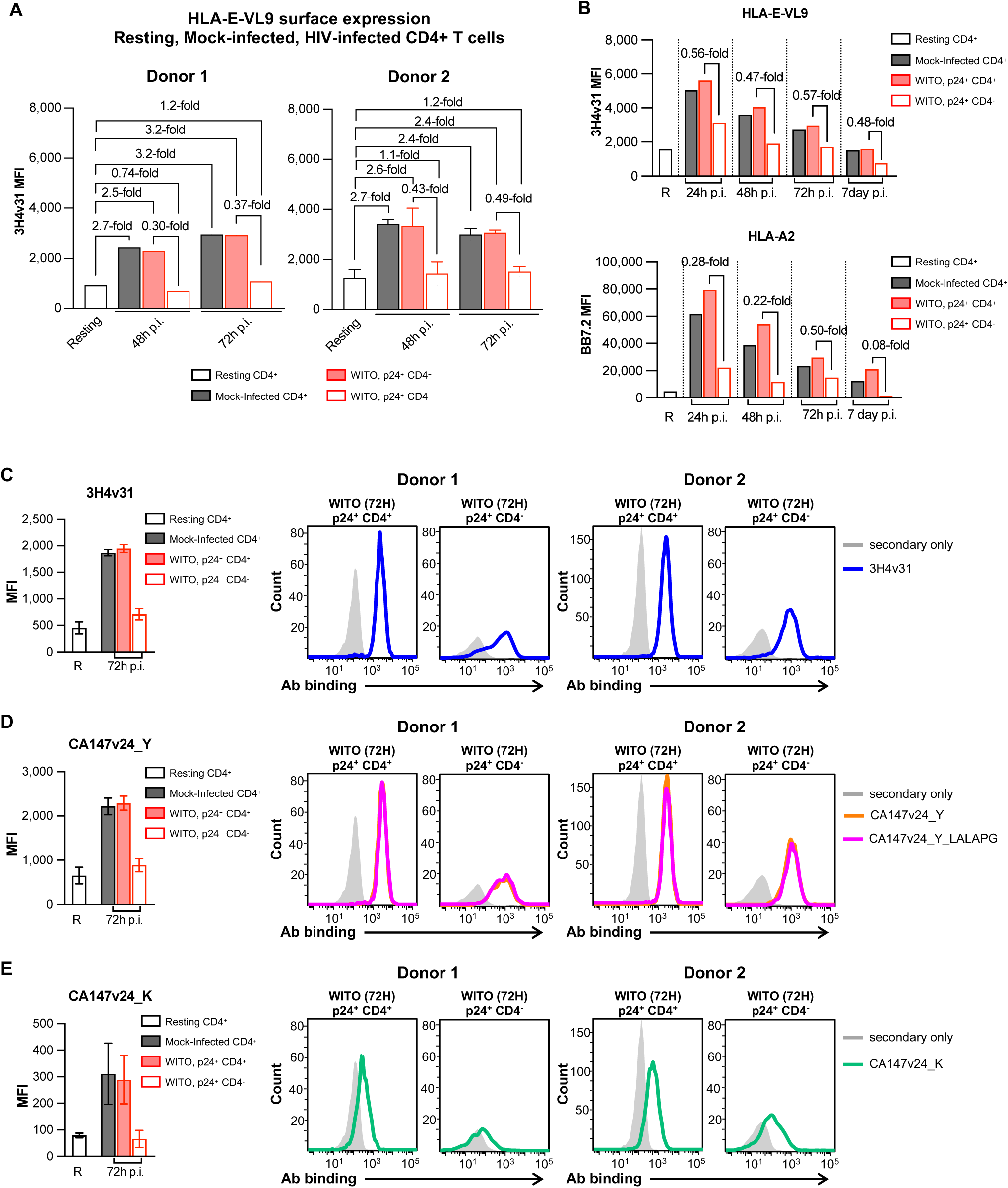
HLA-E-VL9 surface expression on HIV-infected primary CD4+ T cells. Primary human CD4^+^ T cells from two donors were rested overnight, activated for 72 hours, then infected with WITO HIV (WITO) or media (“Mock-infected”) for 72 hours. Donor 1 HLA-E-VL9 expression was assayed at 48 hours post-infection (p.i.), and 72 hours p.i. (one experimental replicate). Donor 2 (HLA-A2 genotype) HLA-E-VL9 expression was assayed concurrently with HLA-A2 surface expression at resting, 24 hours p.i., 48 hours p.i., 72 hours p.i. in one replicate and at resting, 48 hours p.i. and 72 hours p.i. in a second replicate. The two replicates of HLA-E-VL9 surface expression data for donor 2 are shown as mean and standard error of the mean for the resting, 48 hour p.i and 72 hour p.i. timepoints, juxtaposed with HLA-E-VL9 surface expression for donor 1 in panel A, and as separate graphs juxtaposed with HLA-A2 surface expression measured in parallel in (B) and Supplementary Figure 5B. **(A)** Binding of 3H4v31(mouse Fc) to resting, mock-infected, and HIV-infected cells derived from two donors. MFI is displayed after subtraction of background MFI of isotype control mAb. **(B)** Binding of 3H4v31 (mouse Fc) measured concurrently with HLA-A2 on resting, mock-infected, and infected CD4+ T cells derived from donor 2. A second replicate is shown in Figure S5B. **(C-E)** MFI of anti-HLA-E-VL9 mAb 3H4v31 (human Fc) (C), CA147v24_Y (human Fc) (D), CA147v24_K (human Fc) (E) binding to resting, mock-infected and WITO HIV infected cells are shown as bar charts on the left of the panels as bar charts representing mean and SEM of data from two donors. MFI of secondary-only staining controls has been subtracted from the bars. Histograms of binding on the p24^+^ CD4^+^ and p24^+^ CD4^-^ subsets of infected cells on each of two donors is shown to the right of each panel.

Having demonstrated the expression of HLA-E-VL9 on HIV-infected cells using the mouse HLA-E-VL9 mAb 3H4v31, we tested the ability of our human-derived anti-HLA-E-VL9 mAbs CA147v24_Y and CA147v24_K to bind HIV-1-infected primary CD4^+^ T cells (Fig 4 C-E). These anti-HLA-E mAbs mirrored the patterns of 3H4v31 staining of resting, mock-infected, and p24^+^ CD4^+^ and p24^+^ CD4^-^ subsets of HIV-infected cells **(Figure 4C-E).** CA147v24_Y and CA147v24_Y_LALAPG bound similarly, consistent with their Fab affinities **(Figure S5C**).

### Anti-HLA-E-VL9 mAbs trigger NK ADCC of HIV-infected primary CD4+ T cells

We next asked if anti-HLA-E-VL9 antibodies could enhance direct killing and/or trigger primary NK ADCC. Here we have used an infected cell elimination assay in which primary NK cells were co-cultured with autologous activated primary CD4+ T cells infected with the WITO HIV-1 infectious molecular clone 72 hours previously. Antibody-dependent NK killing was quantified as the reduction of live target cells in the presence of test antibodies relative to control wells without antibody (**Figure S5D**).

All anti-HLA-E-VL9 mAbs mediated NK killing of infected cells, tested with primary CD4+ T cell targets isolated from two HIV-seronegative donors and autologous primary NK effectors (**Figure 5A; Figures S5E-F**). Enhancement of NK cell-mediated killing of infected cells occurred mainly via ADCC, as a LALAPG Fc variant of CA147v24_Y (CA147v24_Y_LALAPG), that we had demonstrated produces similar binding patterns to CA147v24_Y (**Figure S5C**), mediated lower levels of killing compared to the full CA147v24_Y antibody (**Figure 5A; Figure S5E**). In relation to p24^+^ CD4^+^ WITO HIV-infected targets, 3H4v31 mediated >90% NK killing at mAb concentrations ≥0.4 μg/mL **(Figure 5A)**. CA147v24_K and CA147v24_Y mediated less killing, consistent with their lower affinity, however, at peak activity, percent cytotoxicity was over 75% against p24^+^ CD4^+^ targets **(Figure 5A).** NK-mediated killing was lower against the p24^+^ CD4^-^ subset of infected cells compared to that against the p24^+^ CD4^-^ subset, in line with lower levels of HLA-E-VL9 on the p24^+^ CD4^-^ cells (**Figure S5A; Figure 4C-E**). Nonetheless, 3H4v31, CA147v24_K, CA147v24_Y, produced as much killing at the positive HIV Ab control, quantified as area under the curve analysis **(Figure S5E)**.

**Figure 5:**
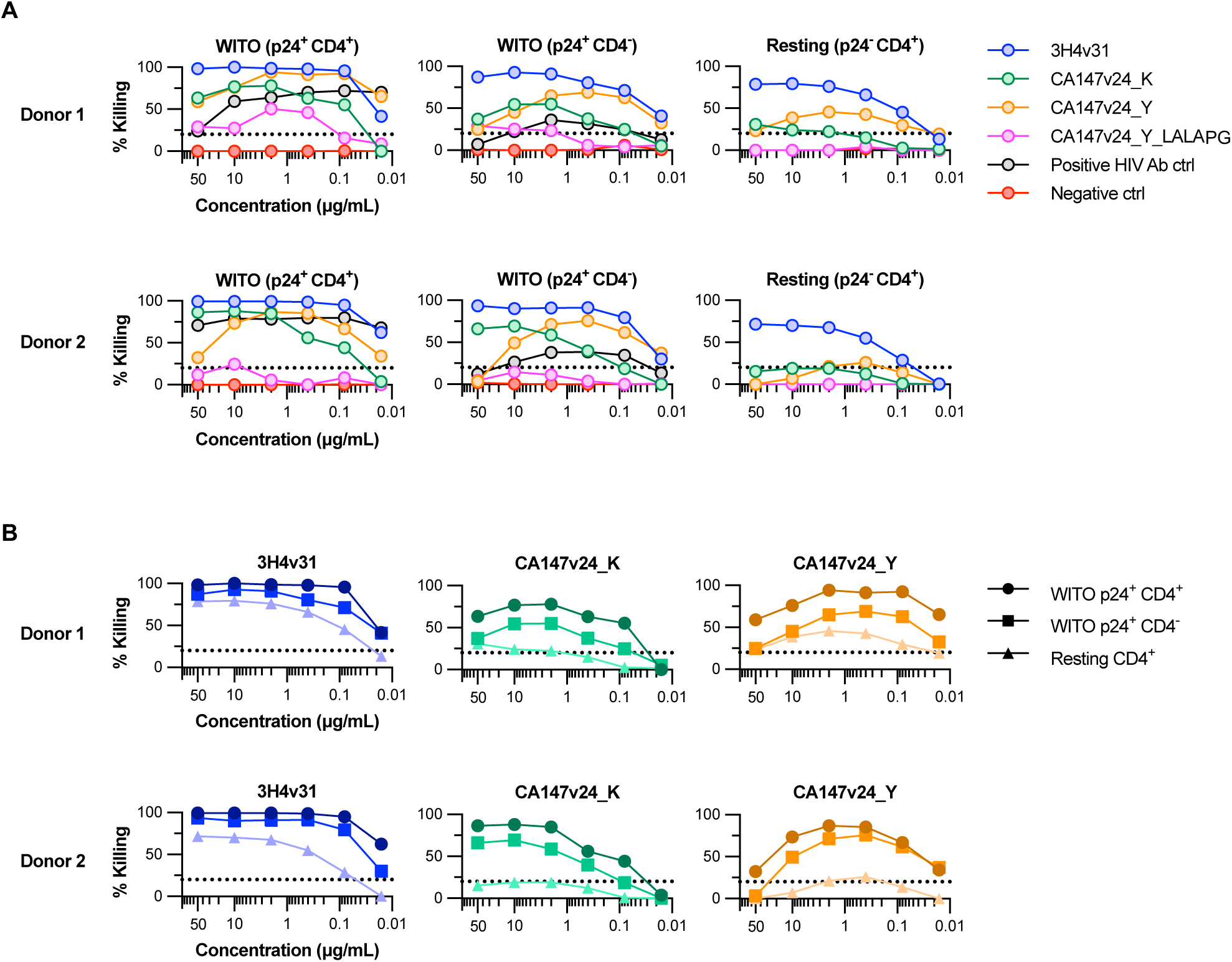
NK ADCC mediated by HLA-E-VL9 mAbs of HIV-infected primary CD4+ T cells. Primary human CD4^+^ T cells from donors 1 and 2 were rested overnight, activated for 72 hours, and Mock- or WITO HIV (WITO)-infected for an additional 72 hours. CD4+ T cells were combined with autologous primary human NK cells for 6 hours with the indicated mAbs at mAb concentrations starting at 50 μg/mL with four subsequent 1:5 dilutions. % Killing was calculated by the reduction of p24^-^ CD4^+^ cells, p24^+^ CD4^+^ cells, or p24^+^ CD4^-^ cells. Previously established positive cutoff for this assay (20% specific killing) is indicated by a dotted line. Killing is displayed in **(A)** overlaying % killing mediated by the indicated antibodies on each of 72-hour WITO infected cells (p24^+^ CD4^+^ subset; left), 72-hour WITO-infected targets (p24^+^ CD4^-^ subset; middle), or resting CD4^+^ cells (right). Positive HIV antibody control in the WITO (p24^+^ CD4^+^) and WITO (p24^+^ CD4^-^) graphs comprises a mix of mAbs A32, 7B2, CH44, and 2G12 that react to HIV-1 gp120 or gp41. The same data shown in A is shown in **(B),** but overlaying % killing on 72-hour WITO infected cells (p24^+^ CD4^+^ subset), 72-hour WITO-infected targets (p24^+^ CD4^-^ subset), and resting CD4^+^ T cells.

All HLA-E-VL9 mAbs mediated less killing of resting, uninfected CD4^+^ T cells than WITO-infected cells (**Figure 5B; Figure S5E**). This preferential NK killing of infected versus resting cells was more pronounced with the lower affinity anti-HLA-E-VL9 mAbs: NK killing of p24^+^ CD4^+^ WITO-infected cells exceeded the killing of resting cells by at least 50% at nearly all tested concentrations between 0.08 μg/mL and 50 μg/mL in the presence of CA147v24_K (K_D_ 290 nM) and CA147v24_Y (140 nM), and only at the lower concentrations of 3H4v31 (K_D_ 62 nM). Higher levels of infected cells compared to resting cells was congruent with the higher levels of HLA-E-VL9 on infected cells; to this point, anti-HLA-E-VL9 mAbs mediated similar levels of NK ADCC of activated mock-infected and the p24^+^ CD4^+^ subset of HIV WITO infected cells **(Figure S5G),** in line with their similar levels of HLA-E-VL9 surface expression (**Figure 4A**). Thus, HLA-E-VL9 mAbs mediated primary NK cell ADCC of HIV-infected CD4^+^ T cells with dose-dependent and affinity-dependent selectivity for activated and/or infected versus resting CD4+ T cells.

### Anti-HLA-E-VL9 mAbs enhance HLA-E-HIV peptide-specific CD8+ T cell activity

Since HLA-E-VL9 is upregulated on activated cells and maintained during the initial stages of HIV infection, we hypothesized that HLA-E-VL9 binding to NKG2A will block TCR-mediated anti-viral activity of NKG2A/CD94-expressing T cells. If so, anti-HLA-E-VL9 antibodies should be able to enhance HLA-E-restricted HIV-specific NKG2A^+^ CD8^+^ T cell killing of HIV infected CD4+ T cells.

To test this, we took advantage of a T cell clone with a TCR specific to a novel HLA-E RevIL9_100-108_ epitope, which we identified computationally as a peptide that can bind HLA-E [52, 53]. This peptide has relatively conservative polymorphism in HIV-1 clade B, with 3 common sequence variants, ILVESPAVL (Rev6), ILGEPPTVL (Rev7) and ILVESPTVL (Rev2B). The 3 variant peptides detectably to HLA-E when tested by sandwich ELISA [52], although binding is relatively weak compared to the VL9 peptide (**Figure S6A**). When expressed as a single-chain peptide-B2m-HLA-E trimer (SCT) in 293T cells the RevIL9 peptide variants also enabled formation of stable cell surface expressed peptide-HLA-E complexes ß2m (**Figure S6B**). The IL9 peptides Rev6 or Rev7 bound to HLA-E-ß2m proteins and formed complexes with moderate thermal stability, as tested by differential scanning fluorimetry (DSF) (**Figure S6C)**. HLA-E tetramer refolded with the IL9 Rev6 peptide was prepared by UV peptide exchange as described previously for HLA-E [53].

PBMCs from a HIV-seronegative HLA-A2-negative blood donor were *in vitro* primed with a pool of three RevIL9 peptide variants (Rev6, Rev7 and Rev2B), using a previously described autologous dendritic cell (DC)-based priming system [34]. A HLA-A2-negative donor was chosen because Rev IL9 is also known to be a HLA-A2 presented epitope. Expansion of IL9-reactive CD8^+^ cells was assessed by staining with HLA-E-Rev6 tetramers (**Figure 6A**). Tetramer-positive CD8^+^ T cells were sorted for single cell cloning. Forty-five RevIL9 specific clones were identified as HLA-E-Rev6 tetramer-positive (median 16.4%, range 3.5% to 51.5%) (**Figure 6B**). T cell receptor sequencing confirmed that each T cell clone was monoclonal with only one TCR*β* chain and one, or rarely two, TCRα chains. Testing with Rev IL9-HLA-E, VL9-HLA-E and IL9-HLA-A2 tetramers confirmed specificity for Rev IL9-HLA-E (**Figure 6C**). A NKG2A/CD94+ RevIL-9 clone, c165, and a NKG2A/CD94- RevIL-9 clone, c186, were further studied (**Figure 6D**).

**Figure 6:**
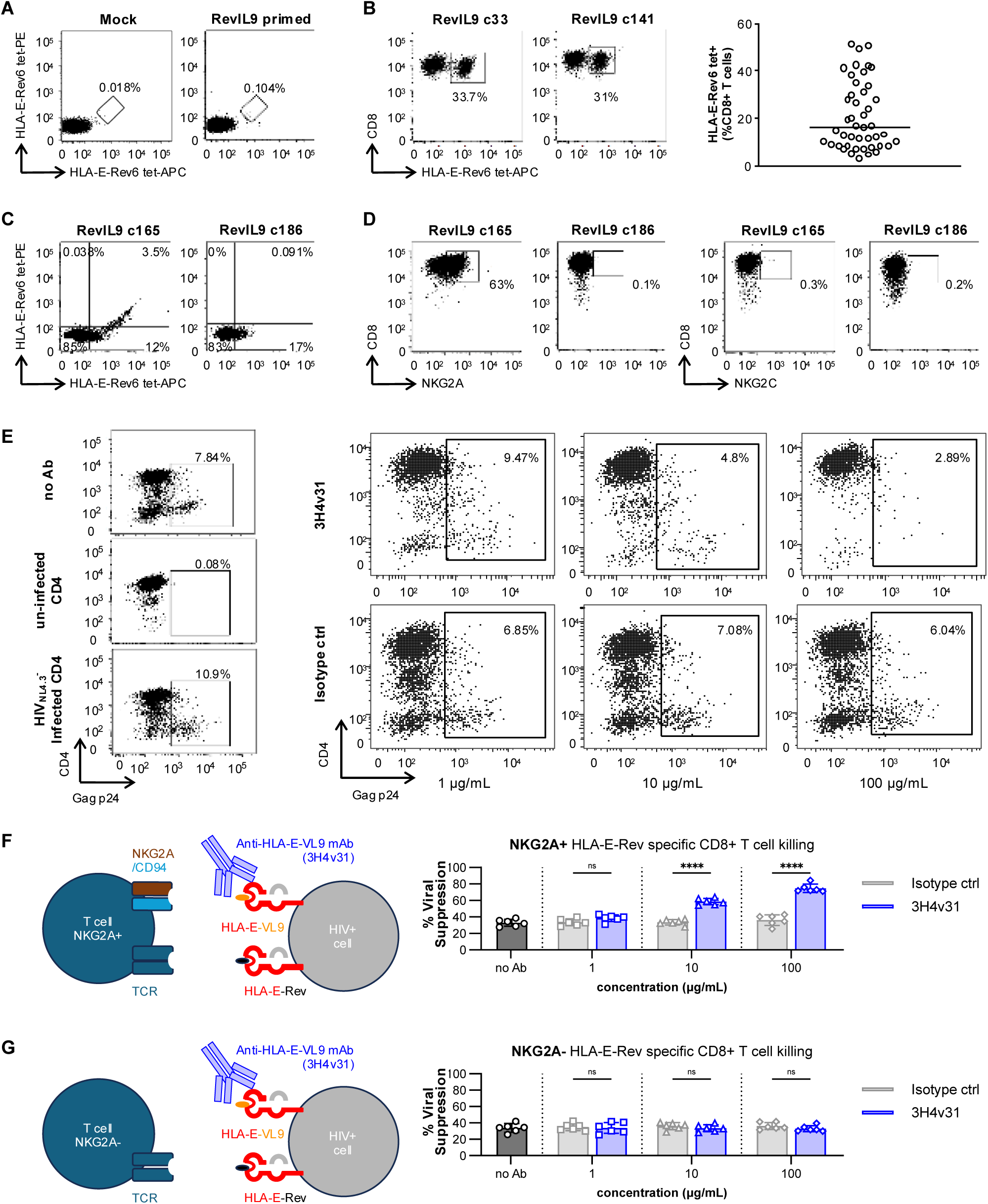
HLA-E-VL9 mAbs enhance cytotoxic CD8^+^ T cell killing. **(A)** Representative FACS plot of co-staining with Streptavidin-APC and PE-conjugated HLA-E-Rev6 tetramers to identify HLA-E restricted Rev6-specific CD8^+^ T cells after mock and the pool of three RevIL9 peptide variants (Rev6, Rev7 and Rev2B) autologous DC priming of PBMC from a HIV-seronegative donor. **(B)** Examples of FACS plots of HLA-E-Rev6 tetramer staining of two RevIL9-reactive CD8+ T cell clones. The frequency of tetramer positive cells for all 45 CD8^+^ T cell clones is also summarized. **(C)** FACS plots illustrate dual positive staining of clone 165 but not clone 186 with HLA-E tetramers refolded with Rev6 and the canonical VL9 signal peptide (conjugated with Streptavidin-APC or Streptavidin-PE respectively). **(D)** Representative FACS plots showing NKG2A and NKG2C staining of clones; c165 expressed NKG2A but not NKG2C. All other clones were NKG2A and NKG2C negative. **(E)** Representative FACS plots of primary human CD4+ T cells infected with HIV-1_NL4.3_ are incubated with HLA-E/RevIL9 specific TCR clones at a 1:1 E:T ratio with HLA-E-VL9 antibody 3H4v31 or isotype control for 5 days and assayed at the end of culture for % remaining infected CD4^+^ T cells as indicated by Gag p24 positivity. **(F)** Enhancement of NKG2A^+^ HLA-E/RevIL9 specific TCR clone killing by 3H4v31 (blue) versus isotype control mAbs (grey). **(G)** No significant enhancement of NKG2A^-^ HLA-E/RevIL9 specific TCR clone killing (p = N.S.). Statistical analysis was performed with two sample unpaired t-test. ****p < 0.0001.

Clonal recognition of naturally presented RevIL9 was evaluated using a previously validated viral suppression assay (VSA), as described previously [34]. CD4^+^ T cells from a HIV-1-seronegative donor were activated, infected with HIV-1 NL4.3 virus, and co-cultured with each clone at an E:T ratio of 3:1 for 5 days. The % inhibition of HIV-1 replication by RevIL9-specific clones was significantly higher than that mediated by the irrelevant, negative control clone, namely a HLA-E-restricted SARS-CoV-2-specific CD8^+^ T cell clones [54] and IL9 tetramer-negative clones included as controls (**Figure S6D-E**). RevIL9-specific clones also inhibited the replication of primary viruses produced from HIV-1 infectious molecular clones (**Figure S6F**).

The ability of 3H4v31 to enhance killing of HIV-infected cells by NKG2A/CD94+ RevIL-9 specific CD8+ T cell clone c165 (**Figure 6C,D**) was then tested. Without antibody, clone c165 suppressed HIV-1 NL4.3 replication inefficiently, reducing the % of infected targets by 35-36%; in the presence of 3H4v31, suppression was significantly enhanced compared to that of isotype control antibody-treated cells, with the reduction in the % of infected targets increasing to a mean of 58.6% with 10 μg/mL of antibody and 75.7% with 100 μg/mL of antibody, (**Figure 6E-F**). Enhanced T cell suppression was not observed with a NKG2A^-^ T cell clone c186 targeting the same Rev epitope (**Figure 6G)**. Thus, blockade of the interaction of NKG2A/CD94 with HLA-E-VL9 with an anti-HLA-E-VL9 mAb enhanced HIV-specific killing mediated by CD8^+^ T cells expressing NKG2A.

## DISCUSSION

Here, we used structure-based design and high throughput library screening to engineer high affinity antibodies specific to HLA-E-VL9. We demonstrated three mechanisms by which anti-HLA-E-VL9 mAbs enhanced HLA-E-VL9^+^ HIV-infected cell killing: inhibitory NKG2A checkpoint blockade releasing NK direct killing (**Figures 2, 3**), triggering NK cell-mediated ADCC **(Figure 5**), and inhibitory NKG2A checkpoint blockade releasing CD8^+^ T cell cytotoxicity (**Figure 6**). Thus, anti-HLA-E-VL9 mAbs have the capacity to engage both NKG2A^+^ NK and T cells and/or CD16^+^ NK functional subsets, and via NK-mediated ADCC, convert HLA-E-VL9 from a critical inhibitory signal protecting HIV-1 to a mediator of HIV-infected cell death (**Figure 5**).

The development of anti-HLA-E-VL9 antibodies enabled us to resolve a major question in the HIV literature, whether HLA-E is upregulated (as proposed by [23]) or downregulated (as proposed by [45]) on infected cells. We found that HLA-E-VL9 surface levels were increased as a function of T cell activation, and persisted on HIV-infected CD4+ targets until the later stages of the viral lifecycle when Nef-mediated CD4 and HLA-A/B downmodulation occurs **(Figure 4, Figure S5B**). Even following downregulation of HLA-E-VL9 on the p24^+^ CD4^-^ subset, HLA-E-VL9 was present at sufficient levels to mediate NK cell killing above that observed on resting CD4^+^ T cells (**Figure 5A, Figure S5E**). We used one HIV strain, WITO [48], in our studies; additional studies will be required to elucidate if there may be HIV strain-specific differences in the kinetics of HLA-E-VL9 surface expression.

Our data that anti-HLA-E-VL9 antibodies enhanced killing of HIV^+^ cells by HLA-E/RevIL9 peptide-specific NKG2A^+^ CD8^+^ T cells (**Figure 6**) highlights that both VL9 and viral peptides are displayed by HLA-E on HIV-infected cells and thus inhibitory checkpoint blockade and ADCC could potentially act synergistically. Anti-HLA-E-VL9 antibody-based therapies may counteract the effect of upregulated HLA-E-VL9 on HIV-infected cells, a viral mechanism that limits CD8+ T cell killing via HLA 1a-peptide recognition. Thus anti-HLA-E-VL9 antibody therapies may also act synergistically with HLA-Ia targeted therapies, and could have broad applicability as an immunotherapy in other contexts such as cancers in which HLA class Ia molecules are downregulated leading to T cell escape [55–58] and where HLA-E is overexpressed [10, 11, 15, 59–61].

Affinity appears to play a crucial role in determining the extent of NK mediated ADCC, with the highest affinity antibody 3H4v31 effecting significantly higher levels of killing of uninfected cells compared to CA147v24_K and CA147v24_Y **(Figure 5B)**. Beyond affinity, NK cells utilize multiple cues to differentiate between self and infected cells, with HLA-E-VL9 representing one such cue [62–64]. Ligands associated with self, such as HLA class Ia molecules HLA-A/B/C, likely protect cells against ADCC, since NK cells have inhibitory KIR and LIR receptors that recognize HLA antigens [65–67]. This has important implications for certain tumors and viruses that downregulate HLA class Ia for T cell evasion. Thus, the HLA-E-VL9 antibodies described here may serve to sensitize NK cells to HIV-infected targets. HLA-E-VL9 mAbs may also have the capacity to engage NK and T cells in mediating cytotoxicity against the many tumors that have been reported to express HLA-E [10–16].

Beyond HLA-E-VL9, our structural analysis of engineered antibodies provides a roadmap on generating similar antibodies against HLA-E in complex with pathogen or tumor specific peptides. Structural analysis revealed key contacts between the antibody and the VL9 peptide that are essential for specificity. In contrast, interactions at position 8, formed strongly by CA117v2v8 and weakly by CA147v24_K and CA147v24_Y, facilitated the recognition of various HLA-E/peptide complexes. However, antibodies that engaged with amino acids at positions 1 and 5 of VL9, such as 3H4v31 and CA147v24, were highly specific for HLA-E-VL9. Thus, structure-based libraries, built like those designed here, could diversify paratope residues to engage exposed residues unique to a given HLA-E/peptide complex. Antibodies against HLA-E/pathogen peptides could be used therapeutically to mediate NK ADCC, or combined with additional treatments to decrease the latent pool of HIV^+^ CD4^+^ T cells (e.g., HIV broadly neutralizing antibodies, latency reactivation agents, cytokines and ART) to synergize in the treatment of people living with HIV [68].

Anti-HLA-E-VL9 antibodies could be used alone or in combination with other antibody or cellular therapies, although several challenges must still be addressed before their therapeutic potential is established. It is unknown if anti-HLA-E-VL9 antibodies can activate endogenous NK cells *in vivo*, or if such cells are present in sufficient numbers in the microenvironment of target cells. Anti-HLA-E-VL9 antibodies could be administered in combination with adoptively transferred NK cells, as we have done in these studies. The NK-92 cell line we tested in our *in vivo* studies has FDA IND approval for human clinical trials and in early trials showed good tolerability and clinical effects in melanoma, lung cancer, acute myeloid leukemia, or Hodgkin Lymphoma [69, 70]. Anti-HLA-E-VL9 antibodies may also increase the activity of NK adoptive cell therapies in development such as CAR NK, which are of growing interest for their safety relative to CAR T cells and potential for use as off-the-shelf therapy [71].

In summary, we demonstrate the development of anti-HLA-E-VL9 antibodies with affinities and specificities that can be finely tuned. This allowed us to select for antibodies that enhanced specific killing without being generally cytotoxic. These antibodies enhanced *in vitro* and *in vivo* killing, via enhancement of direct NK killing, NK-mediated ADCC, and release of CD8^+^ T cell killing by NKG2A/CD94^+^ effectors.

## Limitations of the study

*In vivo* testing of anti-HLA-E-VL9 antibodies in relevant animal models remains limited and will be the major focus of future studies. To better assess anti-tumor activity, anti-HLA-E-VL9 should be tested in more physiologically relevant cancer models that have native, functional NK cells and a more complex tumor environment. While we provide a detailed analysis of HIV-infected cell killing in cell culture, our study is limited by the lack of testing in the SHIV or SIV-infected non-human primate model to determine whether HLA-E-VL9 antibodies can act alone or in concert with other therapies capable of reducing the HIV-infected latent pool of cells. Finally, several tissues express HLA-E, and we demonstrated here that uninfected, resting CD4+ T cells can bind anti-HLA-E-VL9 antibodies. Therefore, the safety of these antibodies is a major concern and will be thoroughly evaluated in pharmacokinetic and safety studies.

## MATERIALS AND METHODS

### Design of scFv libraries

For CA117 mAb, fifty-three residues spread across CDR loops (Supplemental Figure 1A) were randomized in groups of three or four, with all possible combinations of amino acids sampled at these sites, giving a total library size of 5.9 x10^5^ scFv variants. For CA147 mAb, forty-seven residues spread across the CDR loops (Supplemental Figure 1B) were randomized in groups of three, with all possible combinations of amino acids sampled at these sites, giving a total library size of 1.4 x10^5^ scFv variants. For 3H4v3 mAb, three residues in the CDRH3 loop (S100^A^, Y100^B^, G100^C^) were randomized with all possible combinations of amino acids sampled at these sites, giving a total library size of 8 x10^4^ scFv variants. For CA147v24 mAb, four separate libraries were designed, each sampling five residues (Supplemental Figure 2B) as follows: Library 1- HC S31, Y100^B^, Y100^C^; LC S31, Y32; Library 2- HC V97, W98, D99; LC S31, Y32; Library 3- HC Y100^A^, Y100^B^; LC Y91, G92, S93; Library 4- HC S52, G53, S54; LC S31, Y32. Residues listed were randomized with all possible combinations of amino acids sampled at these sites, giving a total library size of 3.2 x10^6^ scFv variants. For CA117, CA147, and 3H4v3, library DNA was synthesized on a BioXP 3250 (Codex) system and amplified with High Fidelity Phusion polymerase (New England Biolabs). For CA147v24, library fragments were produced as gBlock library fragments (IDT), assembled using Gibson assembly (New England Biolabs), and amplified using Q5 polymerase (New England Biolabs).

### Screening of yeast surface display libraries

ScFv libraries were displayed on the surface of yeast as previously described. Briefly, *S. cerevisiae* EBY100 cells (ATCC, MYA-4941) were transformed by electroporation with 4ug of pCTcon2 plasmid (Addgene), previously digested with BamHI, NheI, and SalI enzymes (New England Biolabs), with 12ug of scFv DNA. An aliquot of the transformed cells was serially diluted and plated onto selective SDCAA agar plates (Teknova) to measure library size; determined to be ∼1-5x10^7^ clones for each library. Transformants were expanded in SDCAA media (Teknova) supplemented with pen-strep (ThermoFisher) and grown at 30°C while shaking at 225rpm. Surface expression of the scFv on yeast cells was induced by transferring the transformed cells to SGCAA media (Teknova) at the density of 1x10^7^ cells/ml and allowing for 24-36 hours of growth. 1x10^7^ induced cells were washed twice in ice-cold Sorting buffer (PBS pH 7.4 (Corning), 0.2% BSA (Sigma), 0.1mM EDTA (Sigma)) and incubated with PE labeled HLA-E-VL9 tetramers and FITC labeled anti-c-myc antibodies (ICL) for 1 hour at 4C with rocking. The concentration of the tetramers added to the cells for the initial sort was 5-50ug/mL and decreased for subsequent sorts. After the hour incubation, cells were washed twice with Sorting buffer and sorted on a Sony SH800 FACS or BD FACS-Diva. FITC/PE double positive cells were collected and resuspended in SDCAA media with pen-strep for successive rounds of enrichment. FACS data was analyzed using Flowjo_v10.7 software (BD). The scFv encoding DNA was extracted using the Zymoprep yeast plasmid miniprep II kit (Zymo Research) and transformed into NEB5α E. coli (New England Biolabs) for Sanger sequencing (Genewiz) of individual bacterial colonies.

### Antibody production

Heavy and light chain sequences from the scFvs were synthesized and cloned into IgG1 expression plasmids and antibodies were produced as previously described (S2 helix scaffold paper). Briefly, Expi293F cells (ThermoFisher Scientific) were transiently transfected with an equimolar plasmid mixture of heavy and light chain using Expifectamine (Invitrogen). After overnight incubation, enhancers were added as per the manufacturer’s protocol and the cultures were incubated with shaking for five days at 37 C and 5% CO_2_. The cell culture supernatant was centrifuged and filtered with a 0.8-micron filter. The filtered supernatant was incubated with equilibrated Protein A beads (ThermoFisher) for one hour at 4 °C and washed with 20 mM Tris, 350 mM NaCl at pH=7. The antibodies were eluted with a 2.5% Glacial Acetic Acid Elution Buffer and were buffer exchanged into 25 mM Citric Acid, 125 mM NaCl buffer at pH=6. IgG expression was confirmed by reducing SDS-PAGE and quantified by measuring absorbance at 280nmm (Nanodrop 2000).

### Protein production for crystallographic studies

The HLA-E-VL9 complex was purified from bacterial inclusion bodies as described previously [37]. Rosetta 2(DE3) cells (Novagen) were transformed with plasmids encoding for the HLA-E*0103 heavy chain and beta-2 microglobulin. Cells were induced with 1 mM IPTG at an OD600 of 1.00 and harvested after 18 hours of expression at 20° C. Cells were lysed by sonication in buffer composed of PBS with complete EDTA-free protease inhibitor (Roche) and universal nuclease (Pierce). Lysates were cleared by ultracentrifugation and pelleted inclusion bodies were washed twice with detergent buffer (0.5% Triton X-100, 50 mM TRIS pH 8.0, 100 mM NaCl, 0.1% NaN_3_) and once with 50 mM TRIS pH 8.0, 100 mM NaCl. Washed inclusion bodies were denatured in 8 M urea, 50 mM MES pH 6.5, 1% DTT, 100 mM EDTA and denatured beta-2 microglobulin was injected dropwise into refolding buffer composed of 400 mM L-arginine, 100 mM TRIS pH 8.0, 2 mM EDTA, 5 mM reduced glutathione, 0.5 mM oxidized glutathione. After refolding for 1 hour at 4° C, VL9 peptide (Avantor) resuspended in DMSO was added to the refolding mixture to a final concentration of 20 μM. Denatured HLA-E heavy chain was then added to the refolding mixture in a dropwise manner and the complex was allowed to form while stirring at 4°C for 48 hours. Refolded HLA-E-VL9 complex was then filtered through a 0.45 μm filter and purified by size-exclusion chromatography using a Superdex200 column (Cytiva) in 2 mM TRIS pH 8.0, 200 mM NaCl, 0.02% NaN_3_.

Plasmids encoding the 3H4v31 heavy and light chains were transfected into FreeStyle293 cells using polyethyleneimine. The 3H4v31 heavy chain was modified to include an HRV3C cleavage site between the Fab and Fc domains and a small truncation was made to the C_L_ domain to facilitate crystallization [72]. Protein was purified from filtered cell supernatants using Protein A resin (Pierce) and purified IgG was digested into Fab using HRV3C protease.

3H4v31 Fab was mixed with a 1.5-fold molar excess of HLA-E-VL9 and binding was allowed to occur at 4°C for 1 hour. The complex was then purified by size-exclusion chromatography using a Superdex200 column in 2 mM TRIS pH 8.0, 200 mM NaCl, 0.02% NaN_3_.

### X-ray crystallography

The HLA-E-VL9 + 3H4v31 Fab complex was concentrated to 7.80 mg/mL and used to prepare several crystallization screens using an Oryx4 liquid handling device (Douglas Instruments). Crystals were obtained in a mother liquor composed of 20.1% PEG 1500, 2.01% MPD, 0.2 M magnesium sulfate, and 0.1 M sodium acetate pH 5.5. A single crystal was cryoprotected with mother liquor supplemented with 20% glycerol prior to plunge freezing in liquid nitrogen.

Diffraction data were collected at ALS Beamline 5.0.1. Data were indexed and integrated in iMOSFLM [73] before being merged and scaled in AIMLESS [74]. Despite purification of the intact complex, a molecular replacement solution could only be found for the 3H4v31 Fab, potentially due to the low pH of the mother liquor that facilitated crystallization. Molecular replacement was performed using PhaserMR [75], with PDB ID: 7BH8 chains G and H as a search ensemble [37]. The 3H4v31 Fab molecular replacement solution was iteratively refined using Coot, Phenix and ISOLDE [76–78].

### Cell lines

K562 cells expressing a disulfide-trapped single-chain trimer of HLA-E-beta-2-microglobulin-VL9 (K562 ^HLA-E-VL9^) were made from transduction of the MHC-I null lymphoblast cell line K562 (ATCC, CCL-243) with single chain trimer constructs encoding HLA-E*01:03 with the VL9 peptide VMAPRTLVL, as described in [34] and a gift from Xiaoning Xu (Imperial College London). NK-92 cells (ATCC, CRL 2407), NK-92.CD16 cells (ATCC, PTA-6967) [41, 42], SiHa (ATCC, HTB-35), and SU86.86 (ATCC, CRL-1837) were procured from the ATCC.

### Cell culture

K562 cells were cultured in complete media comprised of RPMI 1640 (Gibco, Cat#11875093) with 10% FBS (Cytiva, Cat#SH3039603), and 100 μg/mL Penicillin-Streptomycin (Gibco, Cat# 15140122). NK-92 cells (ATCC, CRL 2407) and NK-92.CD16 cells (ATCC, PTA-6967) [41, 42] were cultured in MyeloCult H5100 media (StemCell Technologies, Cat#05150) supplemented with 1X Minimal Essential Media Non-essential amino acids solution (Gibco, Cat#11140050), 1mM sodium pyruvate (Gibco, Cat#11360070), 10 mM HEPES (Gibco, Cat# 15630080), 50 μg/mL Penicillin-Streptomycin (Gibco, Cat# 15140122), and 100 IU/mL of HumanKine recombinant human interleukin 2 (Proteintech, Cat# HZ-1015). SiHa cells (ATCC, HTB-35) were cultured in DMEM (Gibco, Cat#11965092) supplemented with 10% FBS (Cytiva, Cat#SH3039603). SU86.86 (ATCC, CRL-1837) were cultured in complete media. All cell lines were maintained at 37°C and 5% CO2 in humidified incubators.

### Chromium-51 (^51^Cr) Release assay

Target cells were washed with complete media comprised of RPMI 1640 (Gibco, Cat# 11875093) with 10% FBS (Cytiva, Cat#SH3039603), resuspended at 1x10^7^ cells per mL, and labeled with 250 μCi/mL of ^51^Cr Na_2_^51^CrO_4_ (Revvity, Cat#NEZ030S001MC). Cells were labeled for 1 hour and 20 minutes at 37°C then washed three times with complete media. Labeled cells were mixed with test antibody at 2 μg/mL and effector cells in at least triplicate wells at the desired effector to target (E:T) ratios in 96-well round bottom plates. After 6 hours of co-culture, cells were pelleted by centrifugation. 25 μL of supernatant was added to 150 μL of Ultima Gold LS Cocktail (Sigma, Cat#12352200) in a 96-well plate. Radioactivity was measured using the MicroBeta TriLux 1450 counter (Perkin Elmer). On every plate, ^51^Cr labeled target cells without effector cells were included as a spontaneous release control, and ^51^Cr labeled target cells mixed with 2% Triton X-100 detergent were included as a maximum release control. The % ^51^Cr release of specific lysis was calculated for each well using the following formula: [(Experimental Release – Spontaneous Release)/(Maximum Release – Spontaneous Release)] × 100.

### CD107a degranulation assay

Effector and target cells were washed with complete media comprised of RPMI 1640 (Gibco, Cat#11875093) with 10% FBS (Cytiva, Cat#SH3039603) and co-cultured in 96-well plates. PE-conjugated anti-human CD107a antibody (clone H4A3, Biolegend, Cat#328607) at 10 μg/mL, in the presence of BD GolgiPlug (BD Biosciences, Cat#555029) and BD GolgiStop (BD Biosciences, Cat#554724) protein transport inhibitors at 1:1000 and 4:6000 concentration respectively. The effector:target cell ratio used was 2.5:1 for experiments using effectors NK-92 cells (ATCC, CRL 2407) or NK-92.CD16 cells (ATCC, PTA-6967), and at 5:1 for experiments using primary NK cells isolated from leukapheresis samples. After 6 hours of co-culture at 37°C ad 5% CO_2_, cells were washed with PBS (Gibco, Cat#14190136) then stained with LIVE/DEAD Fixable Near-IR Dead Cell Stain Kit (Invitrogen, Cat#L34975), followed by another wash with PBS and staining with fluorescently conjugated monoclonal antibodies against APC conjugated anti-human CD56 (clone 5.1H11, Biolegend, Cat#362503) for 20 minutes at room temperature. Cells were washed with PBS then fixed in 2% PFA/PBS and run on a BD Fortessa or BD LSR II flow cytometer and analyzed using FlowJo software, version 10. Cells were gated on the basis of size, viability, GFP, CD56, and CD107a expression. The percentage of NK cells that were positive for CD107a was calculated as the percentage of live GFP-negative CD56^+^ cells that were CD107a^+^.

### Human subjects

Leukapheresis samples were collected from human participants by the External Quality Assurance Program Oversight Laboratory (EQAPOL) at Duke University under principal investigator Dr. Tom Denny. These samples were used in this study as exempt EQAPOL repository samples [79]. All human samples obtained in EQAPOL studies were carried out with the informed consent of trial participants and in compliance with Institutional Review Board protocols approved by Duke University Medical Center.

### Flow cytometry analysis of antibody binding to cell lines

SiHa (ATCC, HTB-35), and SU86.86 (ATCC, CRL-1837) cell lines were released from cell culture plates by incubation with 5mM of EDTA (Thermo Scientific, Cat#R1021) in PBS (Gibco, Cat#14190136) for 3 minutes (SiHa) or 10 minutes (SU86.86) followed by two washes with PBS. Cells were then assayed for viability with LIVE/DEAD Fixable Near-IR Dead Cell Stain (Invitrogen, Cat#L34975) at 1:1000 at room temperature for 10 minutes. Following one wash in PBS, cells were incubated with 3H4v31 IgG1 or isotype control CH65 at 4μg/mL at 4°C for 30 minutes followed by two washes with PBS, then incubated with goat anti-human IgG cross-adsorbed secondary antibody in AF647 (Invitrogen, Cat#A-21445) at 4°C for 30 minutes. Following two washes with PBS, cells were fixed in 2% PFA/PBS and run on a BD Fortessa or BD LSR II flow cytometer and analyzed using FlowJo software, version 10.

### Animals

NOD.Cg-Prkdc scid Il2rgtm1Wjl/SzJ (NSG) mice were purchased from The Jackson Laboratory (Strain 005557). All animal experiments were conducted with approved protocols from the Duke University Institutional Animal Care and Use Committee.

### Orthotopic K562 mouse model

NSG mice were subcutaneously implanted with K562^HLA-E-VL9^ cells over the right flank. Treatments comprised of 10^6^ PBS-washed NK-92 cells, 990 mcg of 3H4v31 or CH65 IgG, and 25000 IU of human IL-2 (Proteintech, Cat# HZ-1015), and delivered intratumorally on days indicated in the text. Tumors were measured daily using calipers. Mice were euthanized at a pre-defined tumor size endpoints (2000 mm^3^ for experiments in Figure 3B, 3375 mm^3^ for experiments in Figure 3C-E).

### Immunohistochemistry

Tumors were excised in their entirety from the subcutaneous R flank. They were sectioned with a scalpel to fit into the OCT cassette, embedded in OCT compound (TFM media, General Data), and the cassette placed on dry ice until the compound was frozen. Cassettes were stored at -80°C. Sections (5 µm) were cut on a cryostat (Thermo Scientific CryoStar NX50) and placed on positively-charged glass slides (Fisher Scientific) and stored at -80C. Prior to antibody incubation, slides were warmed to room temperature and then fixed in 4% formaldehyde (EMS) in fix buffer (10 mM MES pH 6.1, 125 mM KCl, 3 mM MgCl_2_, 2 mM EGTA, 10% Sucrose, all Sigma) for 10 minutes. After washing in PBS, tissue was permeabilized with 0.2% Triton-X-100 (Invitrogen) in PBS for 5 minutes and washed again with PBS. The samples were then incubated with Block-Aid (ThermoFisher Scientific) for 30 minutes and then incubated with primary antibodies diluted in block for 1 hr. Primary antibodies: Goat anti-CD56 (1/400, AF2408, R&D), Rabbit anti-cleaved Caspase-3 (1/400, 9661, Cell Signaling technologies). Samples were washed 3x 5 minutes in PBS and then incubated with labelled secondary antibodies (Donkey anti-Goat Alexa 647, Donkey anti-Rabbit Rhodamine-Red-X, Donkey anti-Human Alexa 790, all Jackson-Immuno) for 45 minutes. Samples were then washed 6 times for 10 minutes in PBS, rinsed with water and mounted under #1.5 coverslips with Prolong Glass (ThermoFisher Scientific).

### Image Acquisition

Slides were imaged using a Nikon 20X λD, 0.8 NA air objective on Nikon Eclipse Ti2 inverted microscope and using Elements acquisition software and LED illumination (Lumencor). Excitation filters: dichroic mirrors: emission filters used were - ET555/20x: T585lxpr: ET595/33m for Rhodamine-Red-X; ET635/30x: T660lpxr: ET697/60m for Alexa 647; FF01-735/28 (Semrock): T760lpxr: ET811/80m for Alexa 790- all filters and dichroics Chroma technology unless noted. Images were obtained using an Orca Fusion sCMOS camera (Hamamatsu). Tumor images were assembled by stitching using Elements software and images were processed in Elements. Regions with high levels of NK-92 cells were selected for comparison of cleaved caspase staining while avoiding any areas that might contain frost damage.

#### Infection of primary cells

Cryopreserved peripheral blood mononuclear cells (PBMCs) were obtained from PBMC from healthy donors obtained from the EQAPOL [79]. PBMCs were thawed and stimulated in R20 media (RPMI media (Invitrogen) with 20% Fetal Bovine Serum (Gemini Bioproducts), 2mM L-glutamine (Invitrogen), 50 U /mL penicillin (Invitrogen), and 50 μg/mL Gentamicin (Invitrogen)) supplemented with IL-2 (30 U/mL, Proleukin), anti-CD3 (25 ng/mL clone OKT-3, Invitrogen) and anti-CD28 (25ng/mL, BD Biosciences) antibodies for 72 hours at 37°C in 5% CO_2_. Primary CD4+ T cells were isolated by CD4+ T cell enrichment according to the manufacturer protocol (CD4+ T cell isolation kit, order no: 130-096-533, Miltenyi Biotech, Germany). 1.5 x 10^6^ cells were infected using 1 mL virus supernatant by spinoculation (1125 x g) for 2 hours at 20 °C. After spinoculation, 2 mL of R20 supplemented with IL-2 was added to each infection and infections were left for 24-72 hours. Cells were plated in 12 well plate at the concentration of 0.5x10^6^ cells/mL in R20+30 U/mL of IL-2.

#### Infected cell antibody binding assay (ICABA)

HIV-1-infected or mock-infected primary CD4+ T cells were obtained as described above. Cells incubated in the absence of virus (mock infected) were used as a negative infection control. Following infection, cells were washed in PBS, dispensed into 96-well V-bottom plates at 2 x 10^5^ cells/well and incubated with 5 μg/mL mAb for 2 hours at 37 °C. Subsequently, cells were washed twice with 250 μL/well of PBS, stained with vital dye (Live/Dead Fixable Aqua Dead Cell Stain, Invitrogen), washed with 1%FBS-PBS; WB and stained with anti-CD4-PerCP-Cy5.5 (clone OKT4; eBiosciences) to a final dilution of 1:40 in the dark for 20 min at RT. Cells were then washed with WB and stained with anti-human secondary APC-conjugated Fab to a final dilution of 1:200 or anti-murine secondary APC-conjugated Fab to a final dilution of 1:500 and incubated in the dark for 25 min at RT. Cells were then washed with WB and resuspended in 100 μL/well Cytofix/Cytoperm (BD Biosciences), incubated in the dark for 20 min at 4 °C, washed in 1x Cytoperm wash solution (BD Biosciences) and stained with anti-p24 antibody (clone KC57-RD1; Beckman Coulter) to a final dilution of 1:100 in the dark for 25 min at 4 °C. Cells were washed three times with Cytoperm wash solution and resuspended in 125 μL PBS-1% paraformaldehyde. The samples were acquired within 24 hours using a BD Fortessa cytometer. A minimum of 50,000 total events was acquired for each analysis. Gates were set to exclude doublets and dead cells. The appropriate compensation beads were used to compensate the spill over signal. Data analysis was performed using FlowJo 10.10.0 software (TreeStar). % Ab binding and MFI from wells which included the secondary antibody alone (no mAb) were subtracted from samples to calculate the %Ab binding or MFI specifically due to mAb binding.

### Natural Killer Cell Infected Cell Elimination Assay

HIV-1 or mock-infected primary CD4^+^ T cells were used as targets. Autologous cryo-preserved PBMCs rested overnight in R10 supplemented with 10 ng/mL of IL-15 (Miltenyi Biotec) were used as a source of effector cells. The day of the assay NK cells were isolated using human NK cell isolation kit (order no. 130-092-657, Miltenyi Biotech, Germany). Infected and uninfected target cells were labelled with a fluorescent target-cell marker (TFL4; OncoImmunin) for 15 min at 37 °C, as specified by manufacturer. Cells were washed in R10 and adjusted to a concentration of 0.2x10^6^ cells/mL. Autologous NK cells were then added to target cells at an effector/target ratio of 5:1 (2 x 10^6^ cells/mL). The target/effector cell suspension was plated in V-bottom 96-well plates and co-cultured with each individual mAb across a range of concentrations using 5-fold serial dilutions starting at 50 µg/mL. Co-cultures were incubated for 6 hours at 37 °C in 5% CO_2_. Subsequently, cells were washed twice with 250 μL/well of PBS and stained with vital dye (Live/Dead Fixable Aqua Dead Cell Stain, Invitrogen) for 20 min at room temperature. Cell were washed with wash buffer (1%FBS-PBS; WB), and stained with anti-CD4-PerCP-Cy5.5 (clone OKT4; eBiosciences) to a final dilution of 1:40 in the dark for 20 min at RT. Cells were then washed with WB, resuspended in 100 μL/well Cytofix/Cytoperm (BD Biosciences), incubated in the dark for 20 min at 4 °C, washed in 1x Cytoperm wash solution (BD Biosciences) and stained with anti-p24 antibody (clone KC57-RD1; Beckman Coulter) to a final dilution of 1:100 for 20 min at 4 °C. Cells were washed three times with Cytoperm wash solution and resuspended in 125 μL PBS-1% paraformaldehyde. The samples were acquired within 24 hours using a BD Fortessa cytometer. A minimum of 2,500 live target cells were acquired for each analysis. Gates were set to exclude doublets and dead cells. The appropriate compensation beads were used to compensate the spill over signal. Data analysis was performed using FlowJo 10.10.0 software (TreeStar).

NK killing was calculated by normalizing counts of CD4^+^, p24^+^ CD4^+^, or p24^+^ CD4^-^ target cells as a percentage of total acquired events from each well. The specific killing of p24^+^ cells was calculated as:

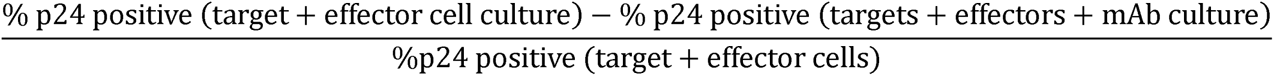

To determine the specific killing of CD4+/- populations, the same equation was used except %p24^+^ was replaced by %p24^+^CD4^+/-^. Synagis (an anti-respiratory syncytial virus monoclonal antibody) was used as a negative control. A combination of four mAbs (A32, 7B2, CH44, 2G12) targeting HIV-1 envelope, was used as a positive control. Assays were conducted once for donor 1 and repeated twice for donor 2. 20% killing was previously determined to be a positive cutoff for ADCC detection by this assay [80].

### Peptides

Synthetic 9 amino acid RevIL9 peptides, ILVESPAVL (Rev6), ILGEPPTVL (Rev7), ILVESPTVL (Rev2B) and HLA-B leader sequence peptide VMAPRTVLL (VL9), were synthesized by Genscript (>85% purity). A UV-labile HLA-B leader-based peptide (VMAPRTLVL) incorporating a 3-amino-3-(2-nitrophenyl)-propionic acid residue substitution at position 5 (J residue) named as 7MT2 was synthesized by Dris Elatmioui at Leiden University Medical Centre, The Netherlands.

### HLA-E binding peptide-exchange ELISA assay

A sensitive HLA-E binding ELISA assay was performed as described previously [34]. Briefly, refolded HLA-E proteins preloaded with a labile VL9 variant peptide (7MT2) was incubated in the presence of excess RevIL9 peptides overnight for exchange reaction. The reaction was added to ELISA plates pre-coated with 20μg/mL anti-human HLA-E mAb (3D12, Biolegend) for 1 hour, plates were then washed with PBS/0.05%Tween-20 and incubated with 2μg/mL anti-human β2M horseradish peroxidase (HRP)-conjugated IgG antibodies for 30 mins. Subsequent enhancement reagent (Dako EnVision) added to amplify the HRP signal followed by TMB substrate for development. Absorbance at 450nm was read using a FLUOstar OMEGA reader. Each run comprised VL9 positive control and a peptide-free no-rescue control to normalize the background and to express the binding affinity as %VL9.

### Differential scanning fluorimetry

The thermal stability of no-peptide and peptide-loaded HLA-E was determined by differential scanning fluorimetry (DSF) using Prometheus NT.48 Series instrumentation (Nanotemper). Assay design was based on a previously published method [34, 54]. In brief, HLA-E was incubated with 10M excess peptide for 30 minutes before split between two Prometheus NT.48 Series nanoDSF Grade Standard Capillaries (Nanotemper, Munich, Germany) and transferred into a capillary sample holder. Thermal melt data calling was automatically generated using the analysis software within PR.ThermControl software version 2.1.5.

### Single Chain Trimers

Single chain trimer constructs were made and tested as previously described [52]. Expression relative to the control construct (which encoded the VL9 peptide VMAPRTLLL) was calculated from median fluorescent intensity (MFI) of the transfected cells, corrected by subtracting the MFI of mock transfected cells.

### In vitro priming and cloning of HLA-E restricted RevIL9 specific CD8+ T cells

PBMCs were isolated from HIV-1 naïve HLA-A*0201 negative donor leukapheresis cones, obtained from NHS Blood and Transplant, UK, by density gradient separation. PBMCs were cultured in AIM-V medium (Invitrogen) with a dendritic cell (DC) differentiation cytokine cocktail of GM-CSF (1000U/ml, Miltenyi Biotech Ltd) and IL-4 (500U/ml, Miltenyi Biotech Ltd) for 24 hrs. DC maturation stimuli of TNF-α (1000U/ml, R&D Systems), IL-1β (10ng/ml, R&D Systems) and prostaglandin E_2_ (PGE_2_ 1μM, Merck) were added together with a pool of RevIL9 peptides (Rev6, Rev7 and Rev2B at a final concentration of 20μM), IL-7 (5ng/ml, R&D Systems) and IL-15 (5ng/ml, R&D Systems). IL-2 was added at a concentration of 500IU/ml at day 6. HLA-E restricted RevIL9 specific CD8+ T cells were evaluated by UV-peptide exchange HLA-E*01:03-Rev6 tetramer conjugated with -APC or -PE staining on day 9. CD3^+^CD4^-^CD56^-^CD94^-^CD8^+^Tetramer-APC^+^/Tetramer-PE^+^ T cells were live sorted using FACSAria III (BD Biosciences). Sorted cells were seeded into 384-well plates at 0.4 cells per well with irradiated (45 Gy) allogeneic feeder cells (3 healthy donors, 2x10^6^ cells/mL) stimulated with PHA (1μg/mL) and IL-2 (500 U/mL) cultured in complete media (CM) containing RPMI 1640, 10% AB human sera (UK National Blood Service), 1% penicillin/streptomycin, 1% glutamine, 1% sodium pyruvate, 1% non-essential amino acids, 0.1% beta-mercaptoethanol. Tetramer positivity was tested using HLA-E tetramers (APC-conjugated) and anti-CD3-APC-Cy7, anti-CD8-BV421, and Live/Dead Fixable Aqua. T cell clones were further expanded with feeder cells and PHA/IL-2.

### HIV-1 Viral suppression / CD8+ T cell infected cell elimination assay

Enriched CD4+ cells were activated with anti-human CD3 at 100ng/ml (clone OKT3, TONBO Biosciences) and IL-2 (100 IU/ml) for 3 days and then infected with the HIV-1 NL4.3 virus obtained from the Programme EVA Centre for AIDS Reagents (National Institute for Biological Standards and Control (NIBSC), a centre of the Health Protection Agency, UK.) at a multiplicity of infection of 1 × 10^−2^ by spinoculation for 2 hours at 27°C, as described previously [81]. HIV-1 NL4.3-infected target cells were cultured in triplicate (1 × 10^5^ cells/well) in RPMI 1640 CM supplemented with 5% AB serum and IL-2 (50 IU/ml), either alone or with RevIL9 clone cells for 5 days at the Effector: Target (E: T) ratio of 3:1. Two irrelevant HLA-E restricted CD8+ T cell clones specific for peptide 001 from SARS-CoV2 ORF1ab protein [54], together with three CD8+ T cell clones, which stained negative with Rev6 tetramer obtained from the same PBMC donor, were used as controls for RevIL9 specificity. At end of coculture, cells were collected and stained with Live/Dead Fixable Aqua before permeabilizing with BD fix/perm solution for intracellular HIV Gag p24 (Beckman Coulter, UK) staining followed by staining with anti-CD3-APC-Cy7, anti-CD8-BV421, anti-CD4-PerCP-Cy5.5 antibodies. The frequency of infected cells was determined by intracellular staining for Gag p24 Ag. Viral inhibition / infected cell elimination was calculated by normalizing to data obtained with no effectors using the formula: (fraction of Gag^+^ cells in CD4^+^ T-cells cultured alone – fraction of Gag^+^ in CD4^+^ T-cells cultured with CD8^+^ clone cells) / fraction of p24^+^ cells in CD4^+^ T-cells cultured alone × 100%.

### Statistical Analysis

Data were plotted using Prism GraphPad version 9. For ^51^Cr release cytotoxic cell death assays and CD107a based degranulation assays, t-tests were applied to evaluate the indicated comparisons, and a P-value of < 0.05 was considered significant. P-values of < 0.05 are indicated with *, < 0.01 with **, and < 0.001 with ***. Kendall Tau correlations were used to test for a dose-dependent effect of antibody concentration on target cell killing. Mixed effects models were used for *in vivo* mouse experiments to compare tumor growth rates between treatment groups. No adjustments were made to the alpha level for multiple comparisons.

## ACKNOWLEDGEMENTS

We thank Duke Human Vaccine Institute (DHVI) programs and finance staff for project oversight and the contributions of technical staff at the DHVI, including Jordan Cocchiaro, Whitney Beck, Cynthia Nagle, Giovanna Hernandez, Esther Lee, Paige Power, Aja Sanzone, Brenna Harrington, Andrew Foulger, Amanda Newman, Pahvie Chhan, Cindy Bowman, Grace Stevens, Thad Gurley, Madison Berry, Kara Anasti, Katayoun Mansouri, Erika Dunford, and Dawn Jones Marshall. We thank Wes Rountree and Yunfei Wang for statistical support. We thank the DHVI Flow Cytometry Core and Duke Cancer Institute Flow Cytometry Shared Resource (FCSR) for technical assistance. We thank Tom Denny for providing leukapheresis samples through EQAPOL (HHSN272201700061C). This study was funded by the Division of AIDS, NIAID, NIH Collaboratory of AIDS Researchers for Eradication (CARE; UM1AI164567) (B.F.H.), the Division of AIDS, NIAID, NIH Consortia for HIV/AIDS Vaccine Development (CHAVD), UM1AI144371 (B.F.H.) and the Bill and Melinda Gates Foundation OPP1108533 (A.J.M.). Support for this study was provided by the Duke Hematology & Transfusion Medicine Training Program (T32 HL007057) to J.K.H. P.B and A.J.M. are Jenner Institute Investigators.

