## Supplementary material for "HLA-E-VL9 antibodies enhance NK cell and CD8^+^ T cell cytotoxicity against HIV-infected CD4^+^ T cells": Table S1

|  | 3H4v31 Fab | CA147v24 + HLA-E-VL9 | CA117v2v8 + HLA-E-VL9 | CA117v2v8 + HLA-E-Mtb44 | CA117v2v8 + HLA-E-RL9HIV |
| --- | --- | --- | --- | --- | --- |
| PDB ID | 9NW5 | 9NW6 | 9NW7 | 9NW8 | 9NW9 |
| *Data collection* | |  | |  |  |
| Space group | *P*2_1_2_1_2_1_ | *P*2_1_2_1_2_1_ | *P*2_1_ | *P*2_1_ | *P*2_1_ |
| Cell dimensions | |  | |  |  |
| a, b, c (Å) | 73.5, 82.8, 94.7 | 73.7, 92.9, 319.3 | 57.5, 71.8, 121.2 | 57.7, 72.3, 121.3 | 57.5, 72.5, 122.4 |
| α, β, γ (°) | 90.0, 90.0, 90.0 | 90.0, 90.0, 90.0 | 90.0, 102.0, 90.0 | 90.0, 101.4, 90.0 | 90.0, 100.8, 90.0 |
| Resolution (Å) | 62.34-2.05 (2.12-2.05) | 79.81-3.10 (3.23-3.10) | 32.23-2.10 (2.17-2.10) | 59.45-2.30 (2.38-2.30) | 40.06-2.30 (2.38-2.30) |
| R_merge_ | 0.823 (1.249) | 0.192 (0.872) | 0.678 (1.688) | 0.744 (1.611) | 0.678 (1.348) |
| I/σI | 12.5 (3.4) | 5.1 (1.7) | 48.9 (2.2) | 9.65 (1.34) | 72.64 (4.07) |
| CC1/2 | 0.940 (0.886) | 0.589 (0.843) | 0.693 (0.396) | 0.567 (0.264) | 0.671 (0.439) |
| Completeness (%) | 99.8 (99.8) | 99.7 (100.0) | 98.4 (98.7) | 93.7 (93.2) | 97.7 (96.7) |
| Redundancy | 6.2 (5.9) | 6.1 (6.3) | 9.1 (9.2) | 8.0 (8.0) | 8.0 (7.7) |
| *Refinement* |  |  |  |  |  |
| R_work_/R_free_ (%) | 21.5/24.4 | 23.9/25.8 | 18.9/23.3 | 19.7/24.7 | 19.1/23.9 |
| No. atoms |  |  |  |  |  |
| Protein | 3339 | 12854 | 6391 | 6516 | 6395 |
| Glycan | 0 | 0 | 0 | 0 | 0 |
| Water | 483 | 0 | 191 | 126 | 121 |
| Average B-factors | |  |  |  |  |
| Protein | 24.4 | 82.1 | 62.9 | 65.6 | 62.9 |
| Solvent | 33.3 |  | 54.7 | 57.6 | 56.2 |
| R.m.s deviations | |  |  |  |  |
| Bond lengths (Å) | 0.005 | 0.011 | 0.011 | 0.009 | 0.011 |
| Bond angles (Å) | 0.77 | 1.29 | 0.99 | 0.89 | 0.94 |
| Ramachandran | |  |  |  |  |
| Favored (%) | 98.4 | 95.8 | 96.9 | 96.7 | 97.2 |
