## Supplementary material for "HLA-E-VL9 antibodies enhance NK cell and CD8^+^ T cell cytotoxicity against HIV-infected CD4^+^ T cells": Table S2

| **HLA-E binding peptides** | | Position | | | | | | | | |
| --- | --- | --- | --- | --- | --- | --- | --- | --- | --- | --- |
|  |  | 1 | 2 | 3 | 4 | 5 | 6 | 7 | 8 | 9 |
| **Self peptides** | VL9 (LVL) | V | M | A | P | R | T | L | V | L |
|  | VL9 (LIL) | V | M | A | P | R | T | L | I | L |
|  | VL9 (VLL) | V | M | A | P | R | T | V | L | L |
| **HIV peptides** | RL9 (HIV) | R | M | Y | S | P | T | S | I | L |
|  | Rev6 (HIV) | I | L | V | E | S | P | A | V | L |
| **Mycobacterium peptides** | Mtb44 | R | L | P | A | K | A | P | L | L |
|  | Mtb14 | R | M | A | A | T | A | Q | V | L |
|  | MtbIL9 | I | M | Y | N | Y | P | A | M | L |
| **SARS-CoV-2 peptides** | SARS-CoV-2-1 | V | M | P | L | S | A | P | T | L |
