## Supplementary figures and images for "HLA-E-VL9 antibodies enhance NK cell and CD8^+^ T cell cytotoxicity against HIV-infected CD4^+^ T cells"

### Figure S1

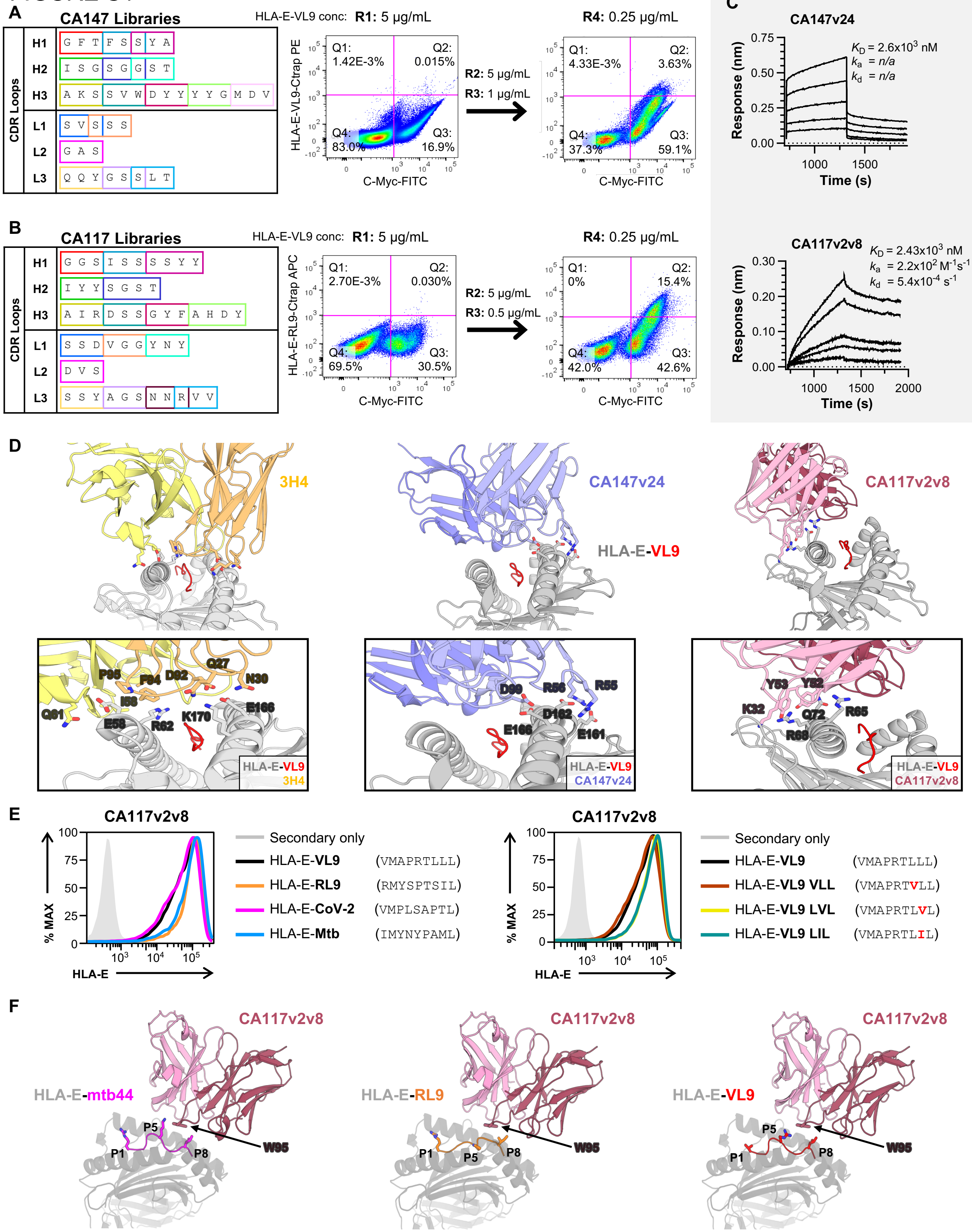

### Figure S2

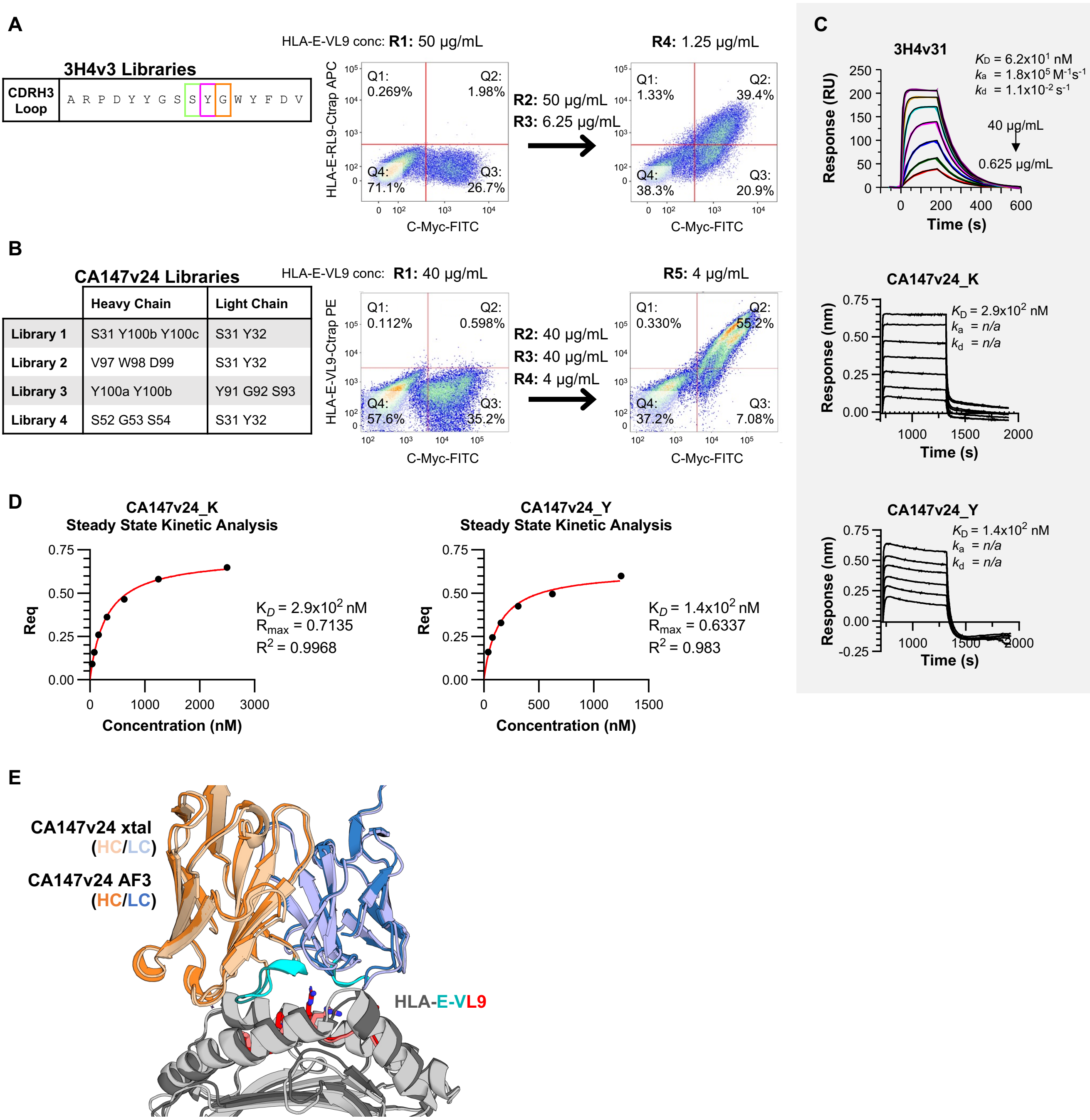

### Figure S3

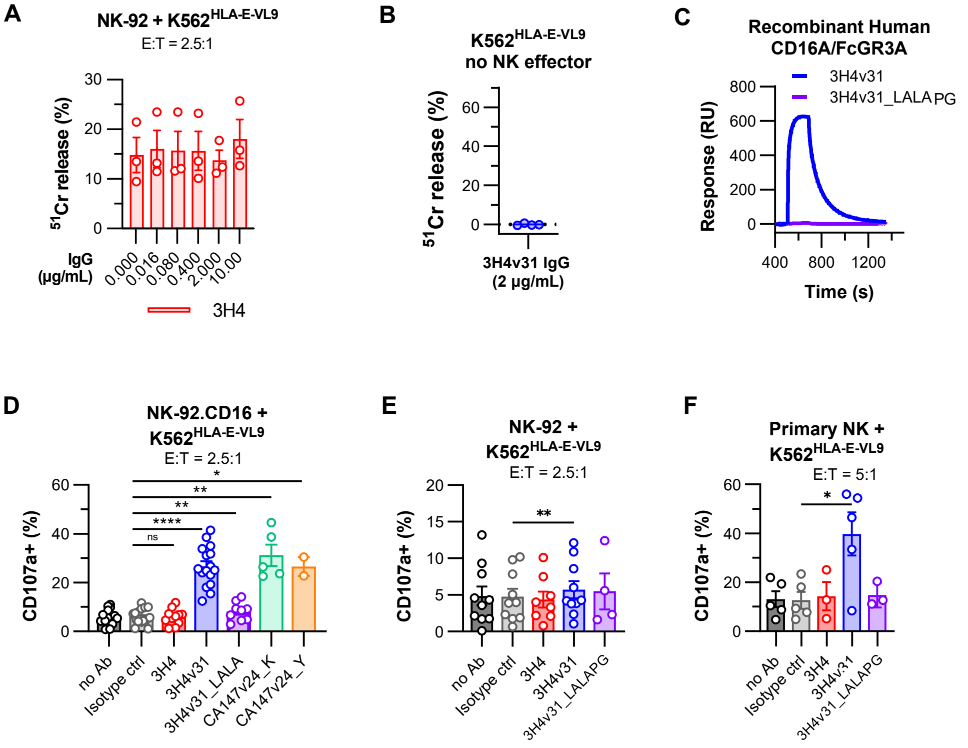

### Figure S5

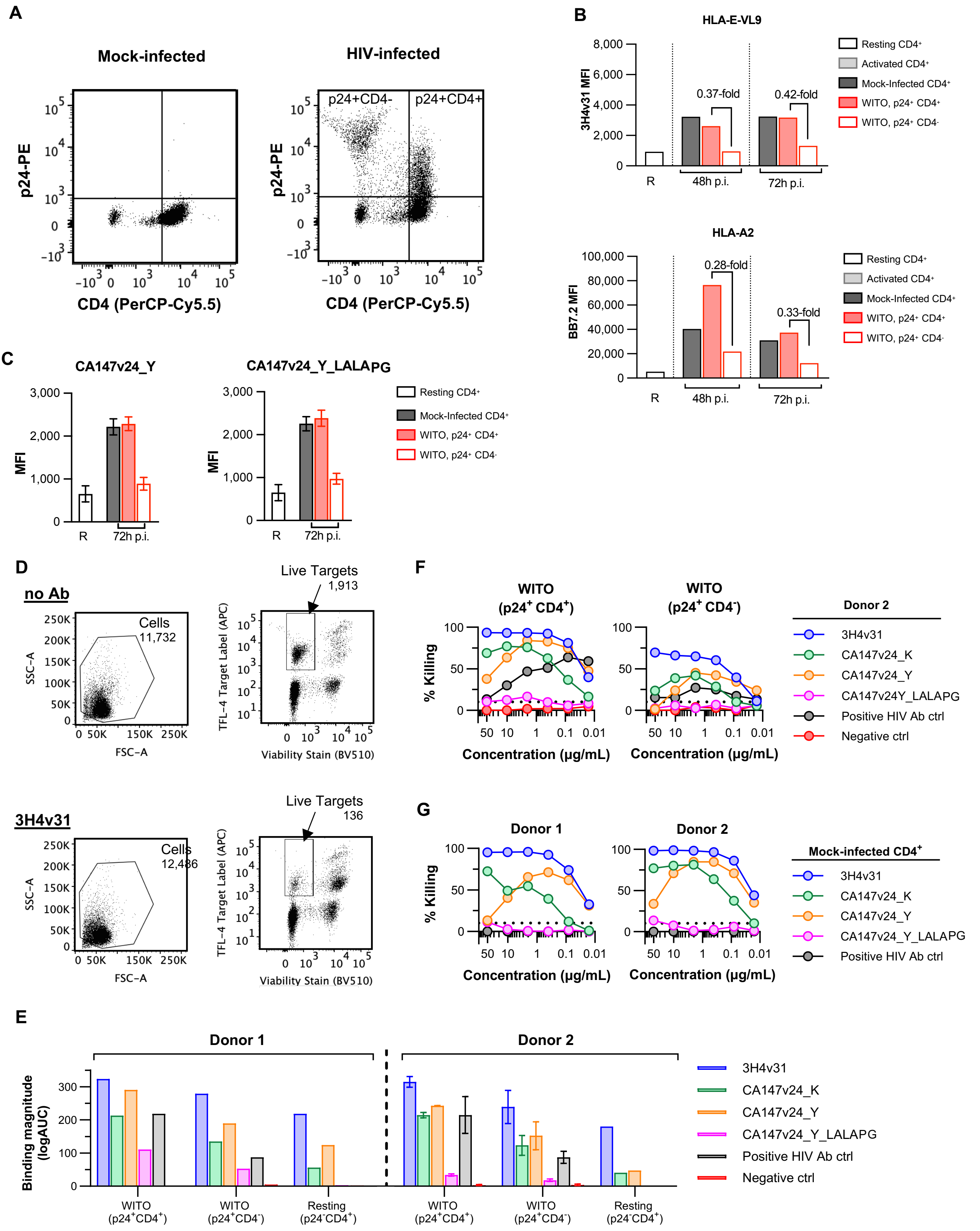

### Figure S6

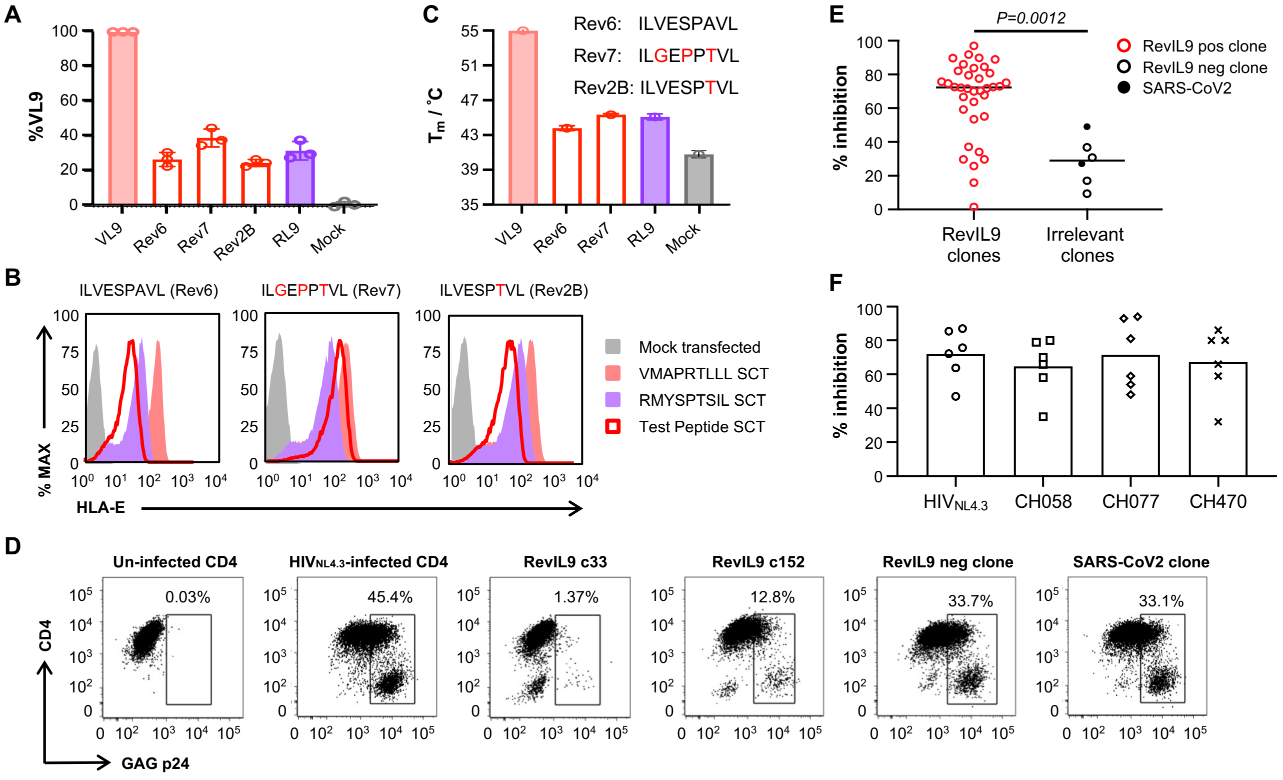
