## Supplementary material for "HLA-E-VL9 antibodies enhance NK cell and CD8^+^ T cell cytotoxicity against HIV-infected CD4^+^ T cells": Figure S4

## A

| Day | 3H4v31 + NK-92.CD16 (mm$^3$) | | | | | | | | | | | Isotype control + NK-92.CD16 (mm3) | | | | | | | | | | | 3H4v31 + NK-92 (mm3) | | | | | | | | | | |
|---|---|---|---|---|---|---|---|---|---|---|---|---|---|---|---|---|---|---|---|---|---|---|---|---|---|---|---|---|---|---|---|---|---|
| | Mus 1 | Mus 2 | Mus 3 | Mus 4 | Mus 5 | Mus 6 | Mus 7 | Mus 8 | Mus 9 | Mus 10 | Mus 11 | Mus 12 | Mus 13 | Mus 14 | Mus 15 | Mus 16 | Mus 17 | Mus 18 | Mus 19 | Mus 20 | Mus 21 | Mus 22 | Mus 23 | Mus 24 | Mus 25 | Mus 26 | Mus 27 | Mus 28 | Mus 29 | Mus 30 | Mus 31 | Mus 32 | Mus 33 |
| **1** | 0 | 0 | 0 | 0 | 0 | 0 | 0 | 0 | 0 | 0 | 0 | 0 | 0 | 0 | 0 | 0 | 0 | 0 | 0 | 0 | 0 | 0 | 0 | 0 | 0 | 0 | 0 | 0 | 0 | 0 | 0 | 0 | 0 |
| **2** | 0 | 0 | 0 | 0 | 0 | 0 | 0 | 0 | 0 | 0 | 0 | 0 | 0 | 0 | 0 | 0 | 0 | 0 | 0 | 0 | 0 | 0 | 0 | 0 | 0 | 0 | 0 | 0 | 0 | 0 | 0 | 0 | 0 |
| **3** | 0 | 0 | 0 | 0 | 0 | 0 | 0 | 0 | 0 | 0 | 0 | 0 | 0 | 0 | 0 | 0 | 0 | 0 | 0 | 0 | 0 | 0 | 0 | 0 | 0 | 0 | 0 | 0 | 0 | 0 | 0 | 0 | 0 |
| **4** | 0 | 0 | 0 | 0 | 0 | 0 | 0 | 0 | 0 | 0 | 0 | 0 | 0 | 0 | 0 | 0 | 0 | 0 | 0 | 0 | 0 | 0 | 0 | 0 | 0 | 0 | 0 | 0 | 0 | 0 | 0 | 0 | 0 |
| **5** | 0 | 0 | 0 | 0 | 0 | 0 | 0 | 0 | 0 | 0 | 0 | 0 | 0 | 0 | 0 | 0 | 0 | 0 | 0 | 0 | 0 | 0 | 0 | 0 | 0 | 0 | 0 | 0 | 0 | 0 | 0 | 0 | 0 |
| **6** | 0 | 0 | 0 | 0 | 0 | 0 | 0 | 0 | 0 | 0 | 0 | 0 | 0 | 0 | 0 | 0 | 0 | 0 | 0 | 0 | 0 | 0 | 0 | 0 | 0 | 0 | 0 | 0 | 0 | 0 | 0 | 0 | 0 |
| **7** | 0 | 79 | 0 | 0 | 0 | 0 | 0 | 0 | 0 | 0 | 0 | 0 | 0 | 0 | 0 | 0 | 0 | 0 | 0 | 0 | 0 | 0 | 0 | 0 | 0 | 0 | 0 | 0 | 0 | 0 | 0 | 0 | 0 |
| **8** | 0 | 122 | 0 | 0 | 0 | 0 | 97 | 0 | 0 | 0 | 0 | 0 | 0 | 0 | 0 | 0 | 191 | 0 | 32 | 0 | 0 | 0 | 0 | 0 | 0 | 0 | 219 | 0 | 107 | 0 | 0 | 0 | 0 |
| **9** | 0 | 257 | 86 | 0 | 279 | 110 | 333 | 0 | 0 | 287 | 185 | 0 | 0 | 0 | 396 | 0 | 455 | 0 | 156 | 0 | 341 | 0 | 176 | 260 | 0 | 84 | 592 | 0 | 108 | 71 | 0 | 0 | 195 |
| **10** | 439 | 271 | 104 | 0 | 294 | 109 | 472 | 273 | 0 | 349 | 248 | 540 | 427 | 0 | 554 | 478 | 613 | 0 | 236 | 0 | 484 | 153 | 177 | 385 | 0 | 94 | 672 | 476 | 155 | 77 | 0 | 0 | 222 |
| **11** | 628 | 433 | 204 | 0 | 275 | 84 | 498 | 319 | 0 | 381 | 245 | 1111 | 729 | 0 | 1016 | 636 | 878 | 0 | 405 | 193 | 580 | 163 | 271 | 662 | 0 | 142 | 855 | 550 | 255 | 89 | 0 | 0 | 223 |
| **12** | 898 | 478 | 214 | 0 | 256 | 74 | 560 | 338 | 0 | 350 | 275 | 1206 | 1005 | 0 | 1100 | 953 | 1572 | 108 | 581 | 253 | 680 | 228 | 260 | 771 | 0 | 178 | 922 | 668 | 289 | 85 | 0 | 0 | 208 |
| **13** | 920 | 657 | 219 | 0 | 268 | 54 | 717 | 297 | 0 | 524 | 267 | 1428 | 1530 | 0 | 1577 | 1435 | 1667 | 280 | 642 | 258 | 814 | 371 | 365 | 1227 | 248 | 192 | 1036 | 986 | 310 | 86 | 0 | 0 | 192 |
| **14** | 983 | 464 | 137 | 0 | 257 | 0 | 918 | 285 | 0 | 393 | 138 | 1724 | 1547 | 0 | 1998 | 2129 | 2047 | 284 | 1107 | 337 | 1060 | 490 | 297 | 1157 | 187 | 132 | 1468 | 1086 | 434 | 55 | 0 | 0 | 118 |
| **15** | 1179 | 732 | 203 | 0 | 297 | 0 | 1204 | 366 | 0 | 458 | 177 | 1967 | 1978 | 108 | 2138 | 2323 | 2108 | 357 | 1476 | 442 | 1709 | 790 | 406 | 1161 | 125 | 71 | 1579 | 953 | 597 | 97 | 0 | 0 | 150 |
| **16** | 1264 | 664 | 150 | 0 | 129 | 0 | 1233 | 272 | 0 | 392 | 105 | 2385 | 2149 | 149 | 2490 | 3061 | 2580 | 503 | 1720 | 553 | 2323 | 899 | 448 | 1242 | 106 | 0 | 1987 | 1089 | 683 | 87 | 0 | 0 | 85 |
| **17** | 1482 | 933 | 176 | 0 | 128 | 0 | 1457 | 246 | 0 | 369 | 121 | 2686 | 2461 | 198 | 3196 | 3807 | 2803 | 617 | 1877 | 822 | 2952 | 1390 | 385 | *n/a* | 0 | 0 | 2237 | 1457 | 812 | 0 | 0 | 0 | 0 |
| **18** | 1651 | 1010 | 170 | 0 | 0 | 0 | 1360 | 227 | 0 | 373 | 69 | 3330 | 3390 | 261 | 3699 | 3807 | 3974 | 773 | 1919 | 948 | 2902 | 1523 | 526 | *n/a* | 0 | 0 | 2242 | 1517 | 879 | 0 | 0 | 0 | 0 |
| **19** | 1876 | 1152 | 183 | 0 | 0 | 0 | 959 | 331 | 0 | 592 | 112 | 3330 | 3390 | 341 | 3699 | 3807 | 3974 | 775 | 2340 | 1043 | 3274 | 1551 | 619 | *n/a* | 0 | 0 | 2468 | 1755 | 1008 | 0 | 0 | 0 | 0 |
| **20** | 2062 | 1397 | 228 | 0 | 0 | 0 | 1542 | 295 | 0 | 762 | 149 | 3330 | 3390 | 533 | 3699 | 3807 | 3974 | 971 | 2530 | 1174 | 3173 | 1901 | 631 | *n/a* | 0 | 0 | 2982 | 1924 | 1016 | 0 | 0 | 0 | 0 |
| **21** | 2450 | 1541 | 240 | 0 | 0 | 0 | 1947 | 380 | 0 | 797 | 127 | 3330 | 3390 | 551 | 3699 | 3807 | 3974 | 1258 | 3046 | 1565 | 4853 | 2133 | 769 | *n/a* | 0 | 0 | 3659 | 2729 | 1448 | 0 | 0 | 0 | 0 |
| **22** | 2572 | 1678 | 268 | 0 | 0 | 0 | 2656 | 533 | 0 | 1173 | 156 | 3330 | 3390 | 752 | 3699 | 3807 | 3974 | 1865 | 4045 | 2398 | 4853 | 3115 | 896 | *n/a* | 0 | 0 | 3659 | 3243 | 1767 | 0 | 0 | 0 | 0 |
| **23** | 3033 | 2050 | 344 | 0 | 187 | 0 | 3016 | 707 | 0 | 1317 | 192 | 3330 | 3390 | 824 | 3699 | 3807 | 3974 | 2520 | 4045 | 3103 | 4853 | 3480 | 1204 | *n/a* | 88 | 0 | 3659 | 4071 | 2138 | 0 | 0 | 0 | 0 |
| **24** | 3785 | 2832 | 323 | 0 | 284 | 0 | 3761 | 944 | 0 | 1505 | 283 | 3330 | 3390 | 898 | 3699 | 3807 | 3974 | 3339 | 4045 | 3384 | 4853 | 3480 | 1383 | *n/a* | 112 | 0 | 3659 | 4071 | 2628 | 0 | 0 | 0 | 0 |
| **25** | 3785 | 3164 | 510 | 0 | 392 | 0 | 3761 | 1174 | 0 | 2062 | 379 | 3330 | 3390 | 1134 | 3699 | 3807 | 3974 | 3339 | 4045 | 3384 | 4853 | 3480 | 1616 | *n/a* | 129 | 0 | 3659 | 4071 | 3081 | 0 | 0 | 0 | 0 |
| **26** | 3785 | 3520 | 492 | 0 | 521 | 0 | 3761 | 1474 | 0 | 2356 | 431 | 3330 | 3390 | 1251 | 3699 | 3807 | 3974 | 3339 | 4045 | 3384 | 4853 | 3480 | 2018 | *n/a* | 138 | 0 | 3659 | 4071 | 3546 | 0 | 0 | 0 | 0 |
| **27** | 3785 | 3520 | 525 | 0 | 600 | 0 | 3761 | 1722 | 0 | 2980 | 674 | 3330 | 3390 | 1533 | 3699 | 3807 | 3974 | 3339 | 4045 | 3384 | 4853 | 3480 | 2160 | *n/a* | 152 | 0 | 3659 | 4071 | 3546 | 0 | 0 | 0 | 137 |
| **28** | 3785 | 3520 | 601 | 0 | 792 | 0 | 3761 | 2223 | 0 | 3779 | 783 | 3330 | 3390 | 1864 | 3699 | 3807 | 3974 | 3339 | 4045 | 3384 | 4853 | 3480 | 2270 | *n/a* | 201 | 0 | 3659 | 4071 | 3546 | 0 | 0 | 0 | 160 |

## B

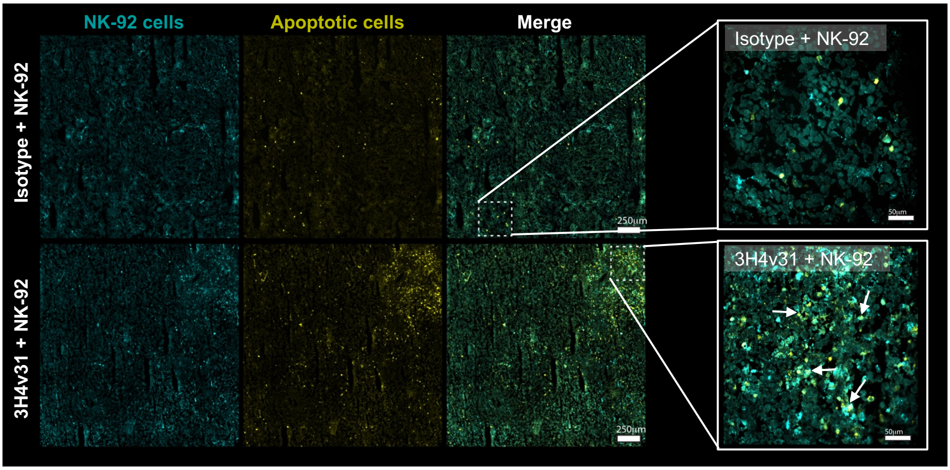


## C

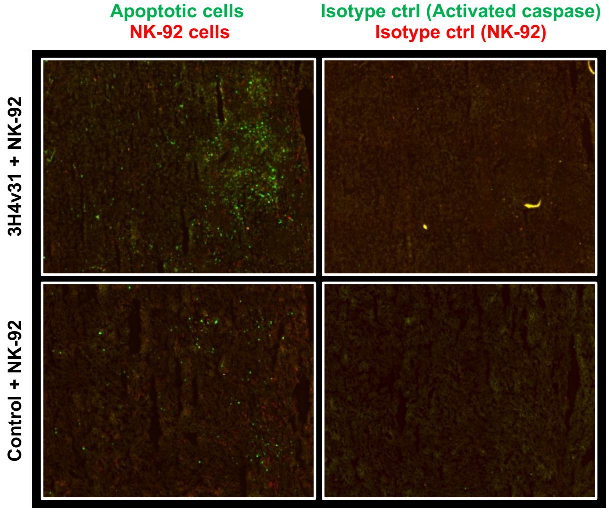
