## Supplemental Figure Legends for "HLA-E-VL9 antibodies enhance NK cell and CD8^+^ T cell cytotoxicity against HIV-infected CD4^+^ T cells"

**Supplementary Figure 1:** **(A, B)** Library design of CA117 (A) and CA147 (B) scFv libraries with groups of randomized residues highlighted in colored boxes (left). Enrichment of HLA-E-VL9 + library clones after four rounds of selection by fluorescence-activated cell sorting (FACS, right). The yeast cells containing the scFv libraries were sorted sequentially for binding to decreasing concentrations of fluorescently labeled HLA-E-VL9 (R1, 5 μg/ml, left; R4, 0.25 μg/ml, right). CA147v24 changes in CDRH2: G55R; S56R; T57E. CA117v2v8 changes in CDRH1 S32K; S33D; Y34V; Y35M and CDRL2 N95W; N95aS: S95bK. **(C)** BLI sensorgrams showing binding kinetics of CA117v2v8 (top) and CA147v24 (lower). For CA117v2v8, rate constants (ka, kd) and dissociation constant K_D_ were determined by curve fitting analysis of SPR data with a 1:1 binding model. For CA147v24, K_D_ was determined by fitting of steady state binding. **(D)** Crystal structures of three different antibodies in complex with HLA-E-VL9 (top). HLA-E heavy chain is shown in *gray*, and the VL9 peptide is in *red*. HLA-E interacting antibody residues are shown in sticks. Detailed view (lower) of the interactions between the antibodies and HLA-E. Antibodies colored as in **(Figure 1C**), HLA-E interacting antibody residues are labeled and shown in sticks. **(E)** FACS binding of CA117 antibodies to HLA-E in complex with either different peptides derived from HIV (RL9), SARS-CoV-2 or *M. tuberculosis* (Mtb) (upper) or with different VL9 variants (lower). (**F**) Crystal structures of CA117v2v8 in complex with HLA-E and 3 different peptides. HLA-E heavy chain is shown in *gray*, MtB44 is *magenta* (left), RL9 is *orange* (middle), VL9 is *red* (right).

**Supplementary Figure 2:** **(A,B)** Library designs of (A) 3H4v3 scFv libraries with randomized residues highlighted in colored boxes and (B) CA147v24 scFv libraries with randomized residues listed (left). Enrichment of HLA-E-VL9 + library clones after four or five rounds of selection by fluorescence-activated cell sorting (FACS, right). The yeast cells containing the scFv libraries were sorted sequentially for binding to decreasing concentrations of fluorescently labeled HLA-E-VL-9 (Rn, X μg/ml denotes concentration of HLA-E-VL9 used “X” for round “n” of selection). CA147v24_K changes in CDRH3: V97I, and CDRL1 loop S31L; Y32M. CA147v24_Y changes in CDRH1: S31W, CDRH3: V97I, and CDRL1 loop S31L; Y32M. 3H3v31 changes in CDRH3: S100aR; Y100bW; G100cN. **(C)** BLI sensorgrams showing binding kinetics of 3H4v31 (top) CA147v24_K (middle), CA147v24_Y (bottom). For 3H4v31, rate constants (ka, kd) and dissociation constant K_D_ were determined by curve fitting analysis of SPR data with a 1:1 binding model. For CA147v24K, K_D_ was determined by fitting of steady state binding. **(D)** BLI steady state binding fit for K_D_ for CA147v24_K and CA147v24_Y. (E) Structural alignment between the experimentally determined structure of CA147v24 in complex with HLA-E-VL9 (*light*) and its model generated by AlphaFold3 (*dark*).

**Supplemental Figure 3: *In vitro* assays of anti-HLA-E-VL9 mAb mediated NK killing.** **(A-B)** Target cell killing was quantified by ^51^Cr release assays. **(A)** Target cell killing by NK-92 with increasing concentration of low affinity 3H4 wildtype version prior to affinity maturation (3H4 WT). n = 3 independent experiments **(B)** Intrinsic cytotoxicity of anti-HLA-E-VL9 was tested for by incubating 3H4v31 IgG with K562^HLA-E-VL9^ cells without any NK effectors. **(C)** Abrogation of binding of the Fc of 3H4v31 by the LALAPG mutation (generating 3H4v31_LALAPG) was tested for by SPR (Biacore T200) assessing binding of 3H4v31 and 3H4v31_LALAPG to CD16/FcγRIII immobilized onto an anti-His/Streptavidin chip. **(D)** Degranulation of NK-92.CD16 effectors upon co-culture with K562^HLA-E-VL9^ and indicated mAbs (n=5 for CA147v24_K, n=2 for CA147v24_Y, n > 8 for other mAb conditions). **(E)**  Degranulation of NK-92 effectors upon co-culture with K562^HLA-E-VL9^ and indicated mAbs (n=4 for 3H4v31_LALAPG, n >7 for other mAb conditions). (**F**) Degranulation of human primary NK cells upon co-culture with K562^HLA-E-VL9^ and indicated mAbs (n=3-5 independent experiments). Dots represent independent experiments and error bars represent standard error of the mean. Statistical analysis by two-sample t-test. *p < 0.05, **p < 0.01, ***p< 0.001, ****p < 0.0001.

**Supplementary Figure 4:** **I*n vivo* assay of anti-HLA-E-VL9 mAb enhancement of NK killing in xenograft mouse model.** **(A)** Table of tumor volume on days 1-28, listed for each of 11 mice per the three treatment groups indicated, in the experiment shown in Main Figure 3. Red text = mice in which tumors progressed on treatment. Green text = mice in which tumors regressed on treatment. **(B)** Immunofluorescence on K562^HLA-E-VL9^ tumors treated with anti-HLA-E-VL9 mAb. Immunofluorescence of fixed K562^HLA-E-VL9^ tumor slices treated with one dose of 3H4v31 + NK-92 cells + IL-2 treatment (*lower panels*) or control mAb CH65 + NK-92 cells + IL-2 (*upper panels*). Tumors were stained for NK-92 cells (*cyan*) and activated caspase as a marker of apoptosis (*yellow*). Detailed view of the merged channels (*right*) highlights areas where NK-92 cells and apoptotic cells are in proximity (*arrows*). **(C)** Immunofluorescence on K562^HLA-E-VL9^ tumors treated with anti-HLA-E-VL9 mAb. Immunofluorescence of fixed K562^HLA-E-VL9^ tumor slices treated with one dose of 3H4v31 + NK-92 cells + IL-2 treatment (*upper panels*) or control mAb CH65 + NK-92 cells + IL-2 (*lower panels*). Tumors were assayed with immunohistochemistry with anti-CD56 antibody for NK-92 cells (*red*) and activated caspase with anti-cleaved Caspase-3 antibody as a marker of apoptosis (*green*). Isotype controls (*right panels)* for NK-92 *(red)* and caspase *(green)* antibodies did not stain the tumors.

**Supplementary Figure 5: HLA-E-VL9 is expressed on HIV-infected CD4^+^ T cells and anti-HLA-E-VL9 mAb mediates NK ADCC**

**(A)** Gating of p24^+^ CD4^+^ versus p24^+^ CD4^-^ subsets of HIV-infected cells. **(B)** Mean fluorescence intensity (MFI) of 3H4v31 binding measured concurrently with HLA-A2 on resting, mock-infected, and infected CD4+ T cells derived from donor 2, second replicate. **(C)** Similar binding to resting, mock- and WITO-infected CD4^+^ T cells by CA147v24_Y and CA147v24Y_LALAPG mAbs. (**D)** Representative flow cytometry histograms showing identification of live and target fluorescent label-4 (TFL-4)-labeled target cells in antibody-mediated NK killing assay. **(E)** Bar charts quantifying area under the curve analysis of killing over concentrations of indicated mAbs shown in Figure 6A. **(F-G)** Killing was calculated by the proportional reduction of TFH-4 labeled CD4+ cells and tested at mAb concentrations starting at 50 μg/mL and four serial 1:5 dilutions . Negative killing values were set to zero. Positive HIV Ab ctrl comprises a mix of mAbs A32, 7B2, CH44, and 2G12 that react with HIV-1 gp120 or gp41. Negative control is anti-respiratory syncytial virus antibody palivizumab. **(F)** NK killing of WITO-infected CD4^+^ T cells derived from donor 2, a second replicate. **(G)** NK killing of mock-infected CD4^+^ T cells derived from donor 1 or donor 2 with indicated antibodies.

**Supplemental Figure 6: HLA-E-VL9 mediates RevIL9-specific CD8+ T cells** **suppression of HIV-1 replication.**

(A-D) Identification of HLA-E binding RevIL9 peptide and *in vitro* suppression of HIV-1 replication by HLA-E restricted RevIL9-specific CD8^+^ T cells. The HIV-1 subtype B RevIL9 peptide and sequence variants that were predicted to bind to HLA-E using NetMHC and an in-house algorithm were experimentally tested for HLA-E binding using 3 approaches (A-C). **(A)** UV peptide-exchange HLA-E binding ELISA assay. The positive control peptide VL9 (VMAPRTVLL) and the previously described HLA-E binding peptide GagRL9 (RMYNPTNIL) were also tested. The average absorbance at 450 nm after subtraction of background of “mock” values is expressed as a % of VL9 binding. The data shown represented three independent experiments. Horizontal lines indicated the mean value. **(B)** Single chain trimer (SCT) HLA-E cell surface expression assay. Histogram overlay plots are representative of at least three repeats. **(C)** Differential scanning fluorimetry (DSF) assay. The thermal melt temperature (Tm) of peptide-free HLA-E–β2m complexes pulsed with 10 M excess of the RevIL9 peptide was shown obtained from three independent experiments. **(D-F)** *In vitro* suppression of HIV-1 replication by HLA-E-RevIL9-reactive T cell clones. CD8 clones (n = 37) were cocultured with HIV-1 NL4.3-infected primary CD4^+^ T cells at an E:T ratio of 3:1. Gag p24^+^ cells were gated on CD3^+^/CD8^-^/CD4^+^ and CD4^-^ cells. HLA-E restricted SARS-CoV2 clones and Rev6 tetramer negative clones obtained from the same PBMC donor were included as negative controls (n = 6). Statistical analysis of the data was performed by a nonparametric Wilcoxon signed rank test. **(F)** *In vitro* suppression of the replication of primary HIV-1 subtype B viruses with differing RevIL9 sequences (HIV-1NL4.3, ILVESPTVL; CH058, VLVESPAVL; CH077, ILVESPTVL; CH470, VLVESPAVL). Horizontal lines indicate the mean of 6 clones tested.
