## Supplemental Table Legends for "HLA-E-VL9 antibodies enhance NK cell and CD8^+^ T cell cytotoxicity against HIV-infected CD4^+^ T cells"

**SUPPLEMENTARY TABLE LEGENDS**

**Supplementary Table 1**: Crystallographic data collection and refinement statistics.

**Supplementary Table 2**: Table listing the sequence of diverse HLA-E binding peptides. Key antibody interacting residues are highlighted in red.
